# Genomic and transcriptomic characterization of Delta SARS-CoV-2 infection in free-ranging white-tailed deer (*Odocoileus virginianus*)

**DOI:** 10.1101/2022.01.20.476458

**Authors:** Jonathon D. Kotwa, Briallen Lobb, Ariane Massé, Marianne Gagnier, Patryk Aftanas, Arinjay Banerjee, Andra Banete, Juliette Blais-Savoie, Jeff Bowman, Tore Buchanan, Hsien-Yao Chee, Peter Kruczkiewicz, Finlay Maguire, Allison J. McGeer, Kuganya Nirmalarajah, Catherine Soos, Lily Yip, L. Robbin Lindsay, Andrew C. Doxey, Oliver Lung, Bradley Pickering, Samira Mubareka

## Abstract

White-tailed deer are susceptible to SARS-CoV-2 and represent a highly important species for surveillance. Nasal swabs and retropharyngeal lymph nodes from white-tailed deer (n=258) collected in November 2021 from Québec, Canada were analyzed for SARS-CoV-2 RNA. We employed viral genomics and transcriptomics to further characterize infection and investigate host response to infection. We detected Delta SARS-CoV-2 (AY.44) in deer from the Estrie region; sequences clustered with human sequences from GISAID collected in October 2021 from Vermont, USA, which borders this region. Mutations in the S-gene and a deletion in ORF8 encoding a truncated protein were detected. Host expression patterns in SARS-CoV-2 infected deer were associated with the innate immune response, including signalling pathways related to anti-viral, pro- and anti-inflammatory signalling, and host damage. Our findings provide preliminary insights of host response to SARS-CoV-2 infection in deer and underscores the importance of ongoing surveillance of key wildlife species for SARS-CoV-2.

## Main text

White-tailed deer (WTD) are considered a highly important species for SARS-CoV-2 surveillance as a result of widespread experimental and epidemiological evidence of SARS-CoV-2 exposure, infection, and transmission across North America ^1–11^. Multiple focally distributed spillover events have been observed, with whole genome sequencing (WGS) analyses initially revealing SARS-CoV-2 lineages that reflect those circulating in humans at the same time ^4,5^. Multiple studies have also documented evidence for onward and sustained deer-to-deer transmission ^1,6,10^. This includes circulation of nearly extinct SARS-CoV-2 lineages (Alpha and Gamma) in WTD in New York and Pennsylvania, USA, with a notable time lapse between detection of these variants in humans and WTD ^1,6^. In Ontario, Canada, we previously identified a highly divergent SARS-CoV-2 variant circulating in WTD (B.1.641) with evidence of deer-to-human transmission ^10^; this finding emphasizes ongoing concerns over the emergence and accumulation of mutations in SARS-CoV-2 while circulating in novel animal hosts ^12^. This introduces the possibility of further divergent evolution and spillback into humans, potentially undermining the effectiveness of medical countermeasures such as antivirals and vaccines ^13–15^.

The growing evidence of SARS-CoV-2 circulation among WTD in North America is suggestive of a new non-human maintenance population or reservoir for the virus. Despite this fundamental change to the ecology of SARS-CoV-2, there is still a dearth of knowledge about the course of infection in WTD, the host-immune response, and resultant evolutionary pressures. For a species to be a competent maintenance host or viral reservoir, the virus needs to persist in a population. One way to achieve this would be for the host to develop immune tolerance against a virus, facilitating virus persistence. For example, bats are broadly considered to be important reservoir hosts for several viruses and are capable of harbouring high viral loads with minimal inflammation and pathology. This is attributed to a balance between immune tolerance (e.g, dampened Stimulator of Interferon Genes (STING) and inflammasome pathways) and host-defence response (e.g., constitutive expression of interferons (IFNs) and interferon stimulating genes [ISGs]) ^16–19^. Alternatively, persistence in the population could be achieved through transmission to naive individuals regardless of host-immunity. There is a need to integrate data on host-immune response with epidemiological and ecological insights to understand whether WTD could represent a competent non-human maintenance population or reservoir for SARS-CoV-2 ^20,21^ and to discern the implications of species-adapted viruses for human health.

We investigated SARS-CoV-2 in WTD in southern Québec, Canada as part of a broader pan-Canadian approach to investigate SARS-CoV-2 spillover into wildlife. We employ viral genomics and transcriptomics to further characterise SARS-CoV-2 infection and host response to infection in comparison to human hosts who clearly exhibit inflammation and disease, anticipating a more subclinical immune profile in infected deer.

## Materials and Methods

### Sample collection and study region

White-tailed deer were sampled during the scheduled hunting season after harvesting by licensed hunters. Samples were collected at two big game registration stations in Dunham and Brownsburg, Québec (**Figure S1**) when harvested and cleaned carcasses were presented for registration; carcasses were returned to the hunters after sampling. Thus, no clinical or post-mortem examination was possible. The Brownsburg station included collection of retropharyngeal lymph node (RPLN) tissues for Chronic Wasting Disease surveillance conducted by the Ministère de l’Environnement, de la Lutte contre les changements climatiques, de la Faune et des Parcs (MELCCFP) in free-ranging WTD in Québec in early November 2021 ^22^. We collected nasal swabs in 1 mL universal transport media and retropharyngeal lymph node (RPLN) tissues were collected in dry 2 mL tubes; both sample types were stored at -80°C prior to analysis. Sex, life stage (juvenile or adult) and geographic location of harvest (latitude and longitude) were recorded for each animal.

### RT-PCR screening and detection

RNA extractions and reverse-transcription polymerase chain reaction (RT-PCR) were performed as described previously ^10^. Briefly, two targets were used for SARS-CoV-2 RNA detection: the 5’ untranslated region (UTR) and the envelope (E) gene ^23^. All samples were run in duplicate and samples with cycle thresholds (Ct) <40 for both SARS-CoV-2 targets and armored RNA enterovirus in at least one replicate were considered positive. Positive samples were further analyzed for a human RNase P gene target to rule out potential contamination ^24^.

Original material from positive samples was sent to the Canadian Food Inspection Agency (CFIA) for confirmatory RT-PCR testing for reporting to the World Organization for Animal Health (WOAH) ^25^ as described previously ^10^. Briefly, confirmatory RT-PCR was performed using primers and probe specific for both SARS-CoV-2 E and nucleocapsid (N) genes ^24^. Samples with Ct <36 for both gene targets were reported to WOAH.

Confidence intervals (CI) for SARS-CoV-2 prevalence of infection were estimated using Stata/SE 15.1 (StataCorp, College Station, Texas, USA; http://www.stata.com) with the Agresti-Coull CI method ^26^. Diagnostic data were plotted on a map of southern Québec according to the latitude and longitude of each WTD with human population density ^27^ and deer harvesting density data ^28^. Graphic displays were produced via QGIS 3.28.2 (Quantum GIS Development Team; http://www.qgis.org).

### SARS-CoV-2 amplification and sequencing

SARS-CoV-2 whole genome sequencing was performed at Sunnybrook Research Institute (SRI) and analyzed at CFIA. Following protocols used by Pickering et al. (2022), all RT-PCR-positive samples were independently 149bp paired-end sequenced using ARTICv3 on Illumina MiniSeq (see supplemental methods for more details).

Paired-end Illumina reads for samples 4055, 4204, 4205 and 4249 were analyzed using the nf-core/viralrecon Nextflow workflow (v2.2) ^29–31^ and lineages assigned with Pangolin (v3.1.17) (see supplemental methods for more details).

Phylogenetic analysis was performed with the WTD consensus sequences generated by nf-core/viralrecon and 93 closely related NCBI and GISAID ^32–34^ sequences identified by UShER ^35^ using a dataset of 14,323,766 genomes from GISAID, GenBank, COG-UK and CNCB (2023-03-28; https://genome.ucsc.edu/cgi-bin/hgPhyloPlace). Multiple sequence alignment (MSA) was performed with Nextalign (v2.13.0) ^36^ of the 4 WTD, 91 NCBI, 2 GISAID and Wuhan-Hu-1 (MN908947.3) sequences. A maximum-likelihood tree was inferred using IQ-TREE (v2.2.0.3) ^37,38^ from the Nextalign MSA with Wuhan-Hu-1 reference strain (MN908947.3) as the outgroup. The best-fit substitution model was determined by IQ-TREE ModelFinder ^39^ to be GTR+F+I+I+R5. The IQ-TREE phylogenetic tree was pruned with BioPython (v1.79) ^40^ for visualization with the R ggtree library (v3.2.0) ^41^. Nexclade CLI (v2.13.0) ^36^ was used to identify amino acid substitutions and deletions in the sequences.

### Virus isolation

Virus isolation was performed on RT-PCR-positive nasal swabs in containment level 3 at the University of Toronto. Vero E6 cells were seeded at a concentration of 3x10^5^ cells/well in a six well-plate. The next day, 250 μL of sample with 16 μg/mL TPCK-treated trypsin (New England BioLabs), 2X penicillin and streptomycin and 2X antibiotic-antimycotic (Wisent; https://www.wisentbioproducts.com/en/)was inoculated onto cells. Plates were returned to a 37°C, 5% CO_2_ incubator for 1 hour and rocked every 15 minutes. After 1 hour, the inoculum was removed and replaced with DMEM containing 2% FBS, 6 μg/mL TPCK-treated trypsin, 2X penicillin/streptomycin, and 2X antibiotic-antimycotic. Cells were observed daily under a light microscope for cytopathic effect for 5 days post infection. The RT-PCR assay was used to confirm SARS-CoV-2 isolation from supernatant. Comparison of isolate sequences to their original nasal swab sample was conducted: cDNA was amplified using ARTIC v4 primer pools (https://github.com/artic-network/artic-ncov2019) and variant calling results were generated using the SIGNAL (SARS-CoV-2 Illumina GeNome Assembly Line) pipeline v1.5.0 ^42^; these were then compared to variant calling data for the original samples using this analysis workflow.

### RNA-sequencing of white-tailed deer nasal swabs

Sufficient material was available for two RT-PCR positive nasal swab samples (4055, 4249) for RNA-sequencing (RNA-seq). As such, eight total RNA samples, including two SARS-CoV-2 RT-PCR-positive and six RT-PCR-negative samples, were submitted for RNA sequencing (RNA-seq) at the Donnelly Sequencing Centre at the University of Toronto (http://ccbr.utoronto.ca/donnelly-sequencing-centre). DNase-treated total RNA was quantified using Qubit RNA HS (cat # Q32852, Thermo Fisher Scientific Inc., Waltham, USA) fluorescent chemistry and 5 ng was used to obtain the RNA integrity number (RIN) using the High Sensitivity RNA ScreenTape (cat # 5067-5579, Agilent Technologies Inc., Santa Clara, USA). Lowest RIN was 1.5; median RIN score was 3. RNA-seq libraries were prepared from RNA samples (150ng) using the NEBNext rRNA Depletion Kit v2 (Human/Mouse/Rat) (NEB Cat# E7405) in conjunction with NEBNext Ultra II RNA Library Prep Kit for Illumina (NEB Cat# E7765). To facilitate the design of deer-specific ssDNA probes Custom RNA Depletion primers were designed using the tool https://depletiondesign.neb.com/. These were then substituted for the oligos provided in the kit. Ribosomal RNA-depleted libraries had a mean concentration of 15.2ng/uL. 1uL top stock of each purified final library was run on an Agilent Bioanalyzer dsDNA High Sensitivity chip (cat # 5067-4626, Agilent Technologies Inc., Santa Clara, USA). The libraries were quantified using the Quant-iT dsDNA high-sensitivity (cat # Q33120, Thermo Fisher Scientific Inc., Waltham, USA) and were pooled at equimolar ratios after size-adjustment. The final pool was run on an Agilent Bioanalyzer dsDNA High Sensitivity chip and quantified using NEBNext Library Quant Kit for Illumina (cat # E7630L, New England Biolabs, Ipswich, USA).

The quantified pool was hybridized at a final concentration of 320 pM and sequenced paired end 150bp on the Illumina NovaSeq6000 platform using a SP flowcell at a depth of 100M reads per sample.

### RNA-seq bioinformatic analysis of white-tailed deer nasal swabs

Raw reads were trimmed with *fastp* v0.21.0 ^43^, with a front 9 nucleotide trim for R1 and R2 based on a FastQC v0.11.4 quality inspection. To remove the deer sequences for microbial profiling, we used STAR v2.5.2b ^44^ to create an index of *Odocoileus virginianus texanus* (GCF_002102435.1; also used as a WTD reference in O’Hara et al., 2022) with a sjdbOverhang of 100 and align the fastp-processed paired-end reads to this index. Deer-aligned reads were removed with BBMap v38.96 filterbyname.sh ^46^. The deer-removed samples were run against the PlusPF pre-built database from May, 17, 2021, consisting of bacteria, archaea, viruses, human, protozoa, and fungi, with Kraken 2 v2.1.2 ^47^. Species abundance was computed with Bracken v2.6.0 ^48^. Distance-based clustering on the species and deer samples was done with the pheatmap v1.0.12 package in R v4.1.1 and applied to a dot plot displaying the relative abundance of species within the community.

For the differential gene expression analysis, the fastp-process paired-end reads were run in the mapping-based mode against a *Odocoileus virginianus texanus* (GCF_002102435.1) decoy-aware (genome as the decoy sequence) transcriptome using Salmon v1.4 ^49^ with the validateMappings setting. The quantification files were imported into R (v4.1.1) and a gene mapping file was created with the makeTxDbFromGFF from the DESeq2 v1.34.0 package ^50^.

Transcript quantifications were merged to the gene-level with tximport using the lengthScaledTPM setting. Low count genes with less than 10 counts were pre-filtered out to improve DESeq2 performance as recommended in the DESeq2 vignette. The differential gene expression analysis was done with the DESeq2 function based on the RT-PCR test results as factor levels. Variance-stabilizing transformation (VST) normalized gene expression for genes that were significantly differentially expressed based on the adjusted *p*-value and had a log_2_ fold change greater than 3 were displayed with gene expression scaling (done with the scale function) in a heatmap with the pheatmap package. The plotPCA function from the DESeq2 package was used to plot a PCA of the VST transformed gene expression profiles. Genes identified as significantly (adjusted p-value) up or down-expressed were run on the Database for Annotation, Visualization and Integrated Discovery (DAVID) ^51^ on Oct. 7 2022 to identify functional terms (including Gene Ontology terms) enriched in the significantly up and down-expressed gene set over a white-tailed deer genome (GCF_002102435.1) background. Notably, these genes and associated processes have been defined using human and mouse studies and as such are used as a surrogate to define likely processes in deer.

To compare the deer results with a human cohort, OrthoFinder v2.5.2 ^52^ was run on a WTD (GCF_002102435.1) and human (GRCh38.p13) proteome, with the proteins mapped back to gene names. The log_2_ fold change values from a previous DESeq2 analysis of RNA-seq human COVID-19 infection gene expression ^53^, was compared to the deer DESeq2 results. Briefly, the human RNA-seq transcriptomics dataset is derived from 50 SARS-CoV-2-positive and 13 SARS-CoV-2 negative individuals; samples were collected from a clinical cohort in the Greater Toronto Area between October 2020 and October 2021 ^53^. The SARS-CoV-2 positive individuals include 16 outpatients, 16 hospitalized (non-ICU) patients, and 18 hospitalized ICU patients ^53^. The GOBP_INNATE_IMMUNE_RESPONSE human gene set from the Human Molecular Signatures Database (MSigDB) C5: ontology ^54^ was used to subset the genes investigated and the correlation between the human and deer immune system log_2_ fold change values was calculated with the cor.test function in R v4.1.1.

### IFITM1 comparison within O. virginianus and mammalian IFITM1 phylogenetics

*IFITM1* duplication in other *O. virginianus* genomes (GCA_023699985.2 and GCA_014726795.1) was confirmed via BLASTN searches of those genomes with the two *IFITM1* sequences (NW_018336621.1:109564-110759 and NW_018336621.1:c138855-136683 for LOC110149600 and LOC110149612, respectively) from *O. virginianus texanus* (GCF_002102435.1). Assemblies GCA_000191625.1 and GCA_000191605.1 were not included as they were extremely small and only fragments of each *IFITM1* gene were able to be detected. As for the IFITM1 sequences across mammals, the IFITM1 HomoloGene group 74501, that includes human, chimpanzee, macaque, wolf, and cow IFITM1 sequences was aligned with MUSCLE v3.8.425 ^55^ to other similar proteins in Boreoeutheria, including the *O. virginianus texanus* IFITM1 proteins. A RAxML-pthreads v8.2.12 ^56^ tree with the PROTGAMMAAUTO setting and the MRE-based Bootstrapping criterion was generated from the alignment, with JTT likelihood with empirical base frequencies being the best-scoring amino acid model for the tree. A nucleotide alignment including coding sequences from all proteins in the tree, as well as the other *O. virginianus* IFITM1 coding sequences, was created with MUSCLE v3.8.425 and subsequently trimmed to the best conserved length with codon alignment corrected. A Fixed Effects Likelihood analysis ^57^ on this alignment generated with the Datamonkey Adaptive Evolution Server ^58^ calculated positions with diversifying selection using the default *p*-value threshold of 0.1.

## Results

### Delta variant of concern detected in white-tailed deer in southern Québec

To discern the prevalence of SARS-CoV-2 in WTD in the region, 258 WTD were sampled in two areas from southern Québec, Canada between November 6–8 2021. The majority of the sampled WTD were adult (92%) and were male (79%). We collected 251 nasal swabs and 104 RPLNs and tested for the presence of SARS-CoV-2 RNA by RT-PCR. Longitude and latitude data were obtained for 257 WTD.

Four nasal swabs were RT-PCR-positive, three of which were confirmed by the CFIA and thus reported to the World Organization for Animal Health (WOAH) as the first cases of SARS-CoV-2 identified in Canadian wildlife on December 1, 2021 (**Table S1**) ^59^. Human RNase P was not detected in any of the positive nasal swabs, excluding contamination from human hosts. Of all nasal swabs, 1.6% (4/251; 95% CI 0.5–4.2%) were positive for SARS-CoV-2 RNA (**Table 1**); no RPLNs were positive. All positive deer were adults and three of the four were male. The four SARS-CoV-2-positive deer were harvested through licensed hunting activity in the high deer density region of Estrie (**Figure 1B**). No RPLNs were available from deer with SARS-CoV-2 positive nasal swabs.

**Table 1.**
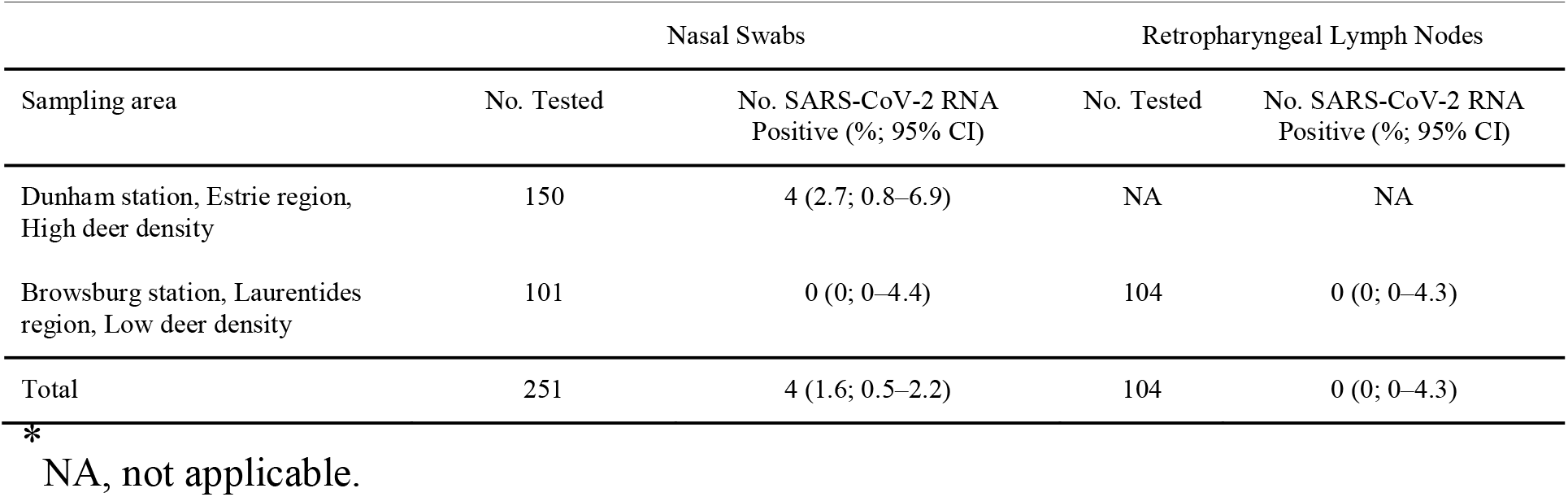
Results from SARS-CoV-2 RT-PCR testing of nasal swabs and retropharyngeal lymph node tissue, and antibody testing of thoracic cavity fluid from white-tailed deer, in two sampling regions, southern Québec, Canada, November 6-8 2021^*^

**Figure 1.**
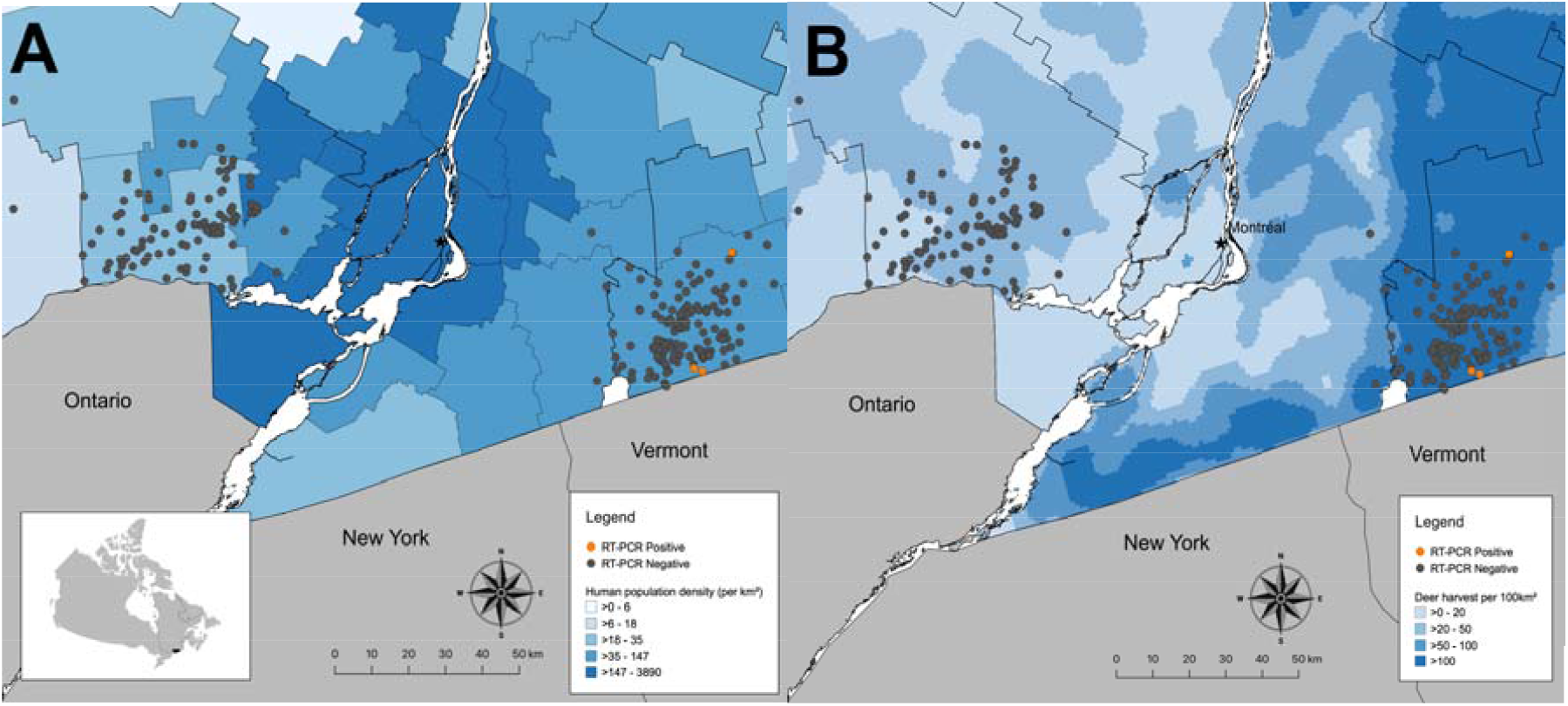
Map of southern Québec with locations of SARS-CoV-2 RT-PCR-positive (orange) and -negative (grey) white-tailed deer from November 6 - 8 2021 superimposed on (A) a choropleth map of human population density (per km^2^) by regional county municipalities (thin grey boundaries) and (B) a heatmap of deer harvest density per 100km^2^ from 2020 as a proxy for deer population density. Inset shows location of Québec (outlined) and study region (shaded black) within Canada.

Whole genome sequencing for SARS-CoV-2 conducted on three confirmed positive samples generated genome coverage of over 95% with equal to or greater than 10X coverage (mean depth from 1096.8 – 1894.8). On the fourth, higher cycle threshold (Ct) sample, 69.1% genome coverage with at least 10X coverage (mean depth of 116.5) was obtained (**Table 2**).

**Table 2.**
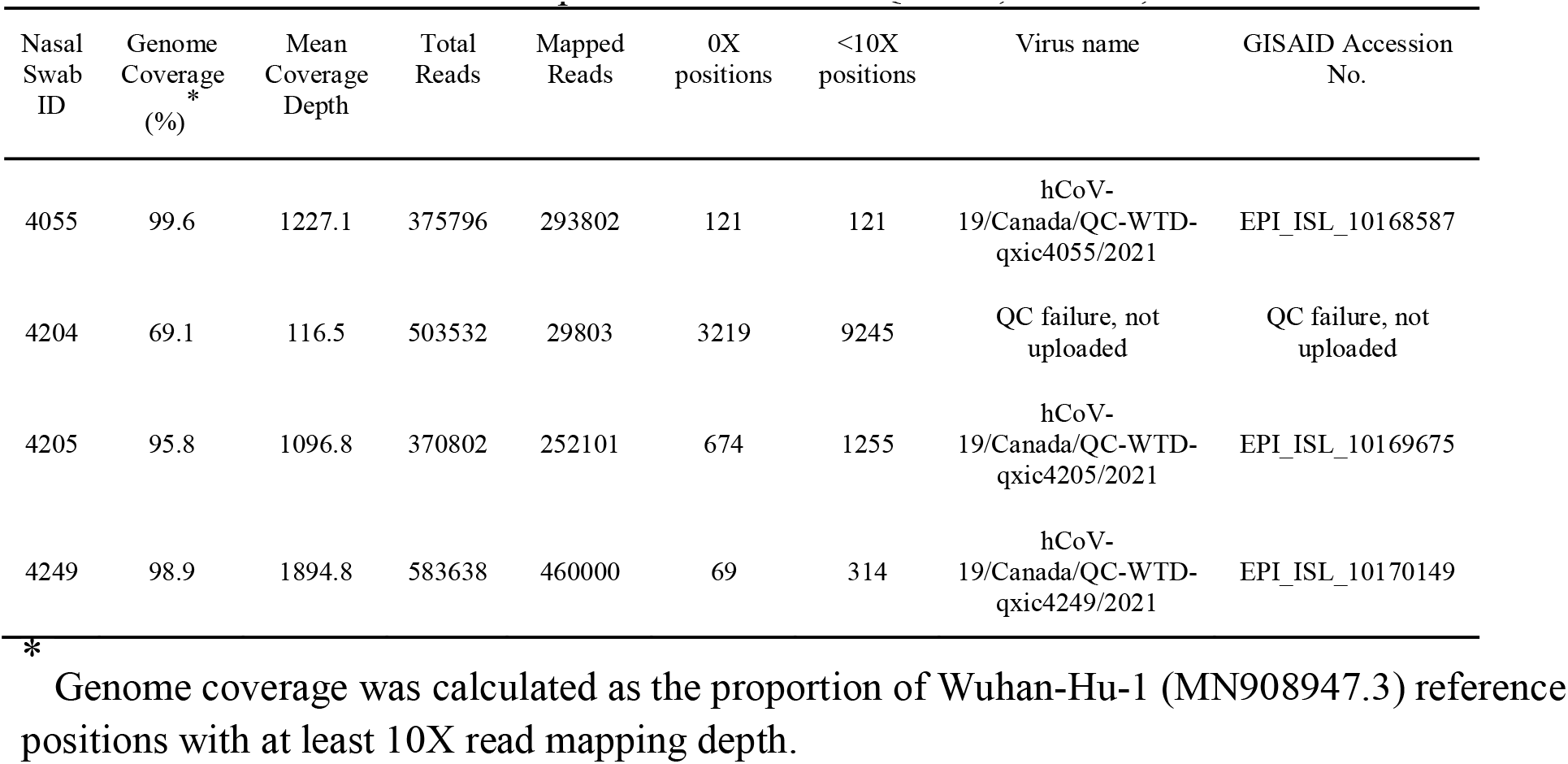
Read mapping statistics from nf-core/viralrecon analysis for four SARS-CoV-2 positive white-tailed deer nasal swab samples from southern Québec, Canada, November 6-8 2021

| Nasal Swab ID | Genome Coverage<sup>*</sup> (%) | Mean Coverage Depth | Total Reads | Mapped Reads | 0X positions | <10X positions | Virus name | GISAID Accession No. |
| --- | --- | --- | --- | --- | --- | --- | --- | --- |
| 4055 | 99.6 | 1227.1 | 375796 | 293802 | 121 | 121 | hCoV-19/Canada/QC-WTD-qxic4055/2021 | EPI_ISL_10168587 |
| 4204 | 69.1 | 116.5 | 503532 | 29803 | 3219 | 9245 | QC failure, not uploaded | QC failure, not uploaded |
| 4205 | 95.8 | 1096.8 | 370802 | 252101 | 674 | 1255 | hCoV-19/Canada/QC-WTD-qxic4205/2021 | EPI_ISL_10169675 |
| 4249 | 98.9 | 1894.8 | 583638 | 460000 | 69 | 314 | hCoV-19/Canada/QC-WTD-qxic4249/2021 | EPI_ISL_10170149 |

\*
Genome coverage was calculated as the proportion of Wuhan-Hu-1 (MN908947.3) reference positions with at least 10X read mapping depth.

Sequences were assigned to lineage AY.44, a sublineage of B.1.617.2 (Delta), with Pangolin (0.96-0.99 ambiguity score), while one sample could not be confidently assigned to a Pangolin lineage due to the large number of N bases (31%) in the consensus sequence. Phylogenetic analysis revealed that all sequenced samples clustered together and shared a most recent common ancestor with SARS-CoV-2 sequences from humans in Vermont, USA between 2021-10-14 and 2021-10-27 (**Figure 2**).

**Figure 2.**
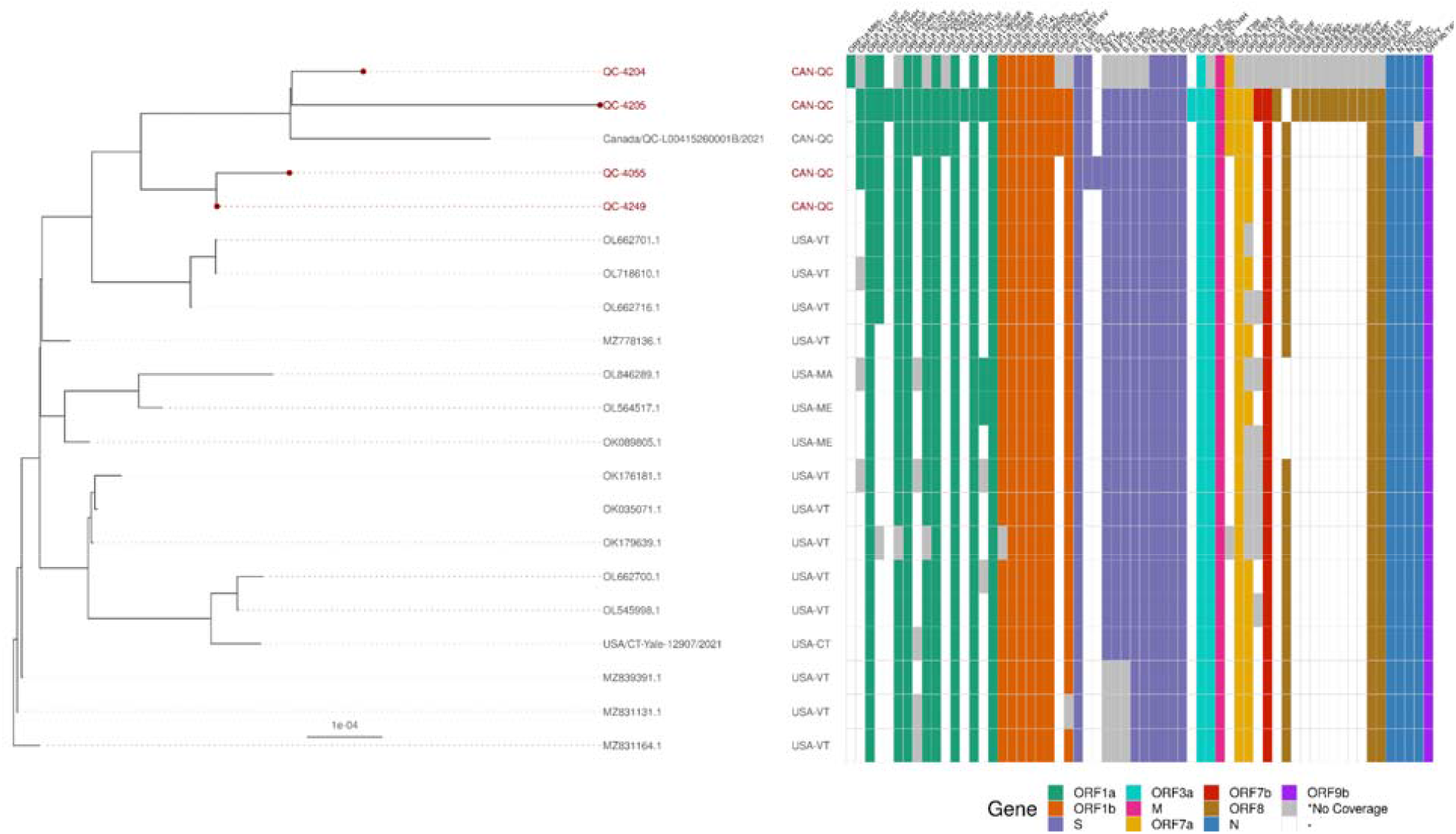
Whole-genome phylogenetic tree of four SARS-CoV-2 positive white-tailed deer sequences and 17 closely related Canadian and American sequences identified by UShER analysis. IQ-TREE inferred the maximum-likelihood phylogenetic tree with a GTR+F+I+I+R5 substitution model (selected by IQ-TREE’s ModelFinder) from a Nextalign multiple sequence alignment of the 4 WTD, 91 NCBI, 2 GISAID and Wuhan-Hu-1 (MN908947.3) sequences. The tree was manually pruned with BioPython to highlight distinct clades and amino acid mutation patterns. The tree was visualized using the ggtree R library. Amino acid (AA) substitutions and deletions in GISAID and white-tailed deer sequences were determined using Nextclade for visualization alongside the tree and other metadata. Some positions within sample 4204 had low or no coverage, however, despite the poor coverage of sample 4204, it still clustered with the other white-tailed deer sequences.

### Mutations in the S gene and ORF8 detected in deer derived SARS-CoV-2 sequences

Mutations were observed in the four positive WTD samples (**Table 3**). Notably, two S gene mutations were observed in the WTD sequences: S:T22I in three samples; S:A27V in one sample only. The S:T22I mutation was observed in only one closely related AY.44 sequence from a human from Quebec while S:A27V was not observed in closely related AY.44 sequences from GISAID. The S:T22I mutation has been observed in 16,628 GISAID sequences as of 2023-04-17 from a multitude of lineages. The S:A27V mutation has been observed in 5,889 GISAID sequences as of 2023-04-17. The S:G142D mutation, which is present in 64% of AY.44 sequences in GISAID (171,978/267,019 sequences as of 2023-04-17), is present in two samples as a minor variant (57% and 53% allele fraction, respectively). The S:G142D mutation is prevalent in many Delta sublineage sequences and present in 10,655,444 GISAID sequences as of 2023-04-17. Interestingly, the S:G1085R mutation, which is present in all four sequenced samples and related AY.44 sequences, is only present in 0.1% (279/267,019) in lineage AY.44 and has been identified in 2,574 GISAID sequences as of 2023-04-17.

**Table 3.**
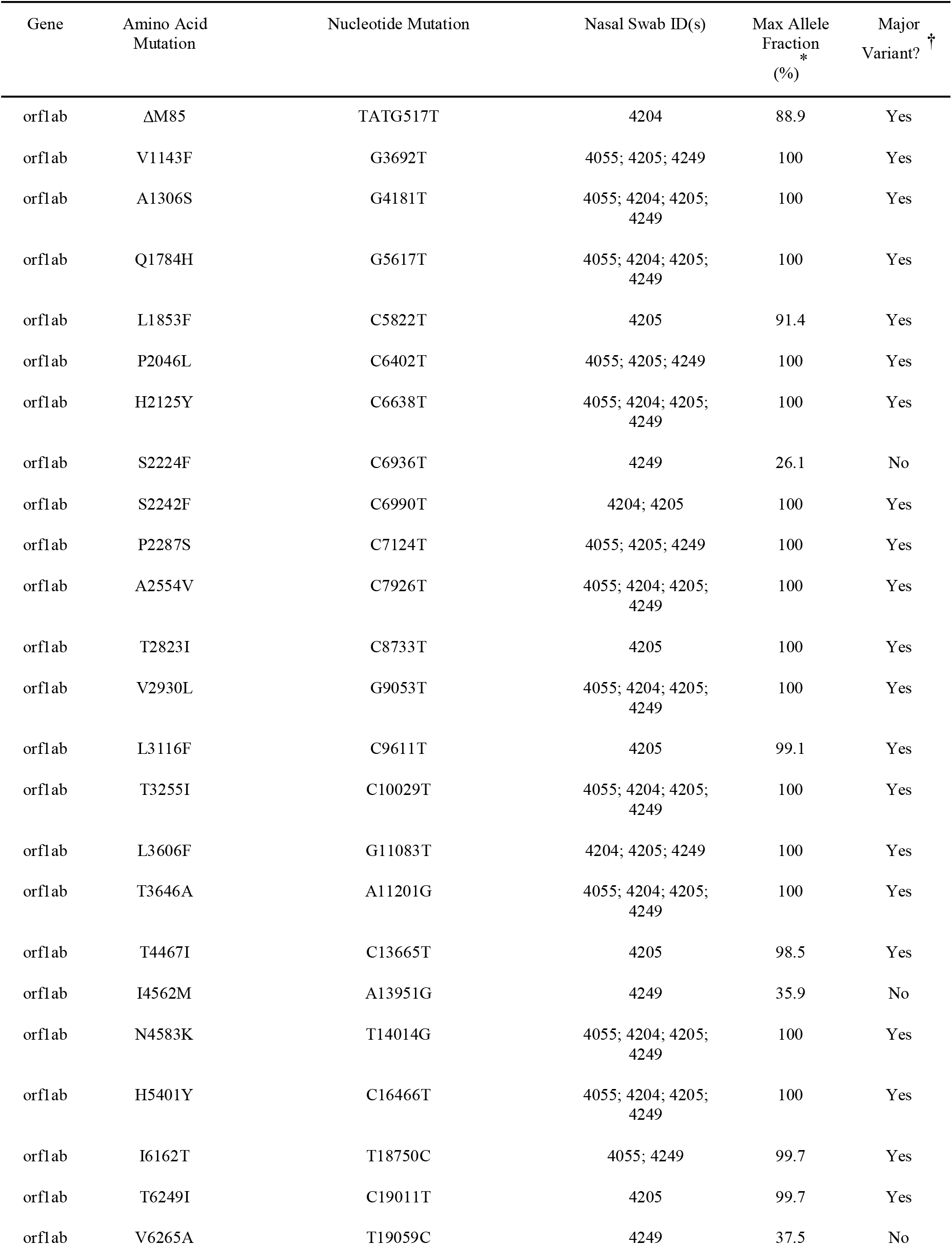
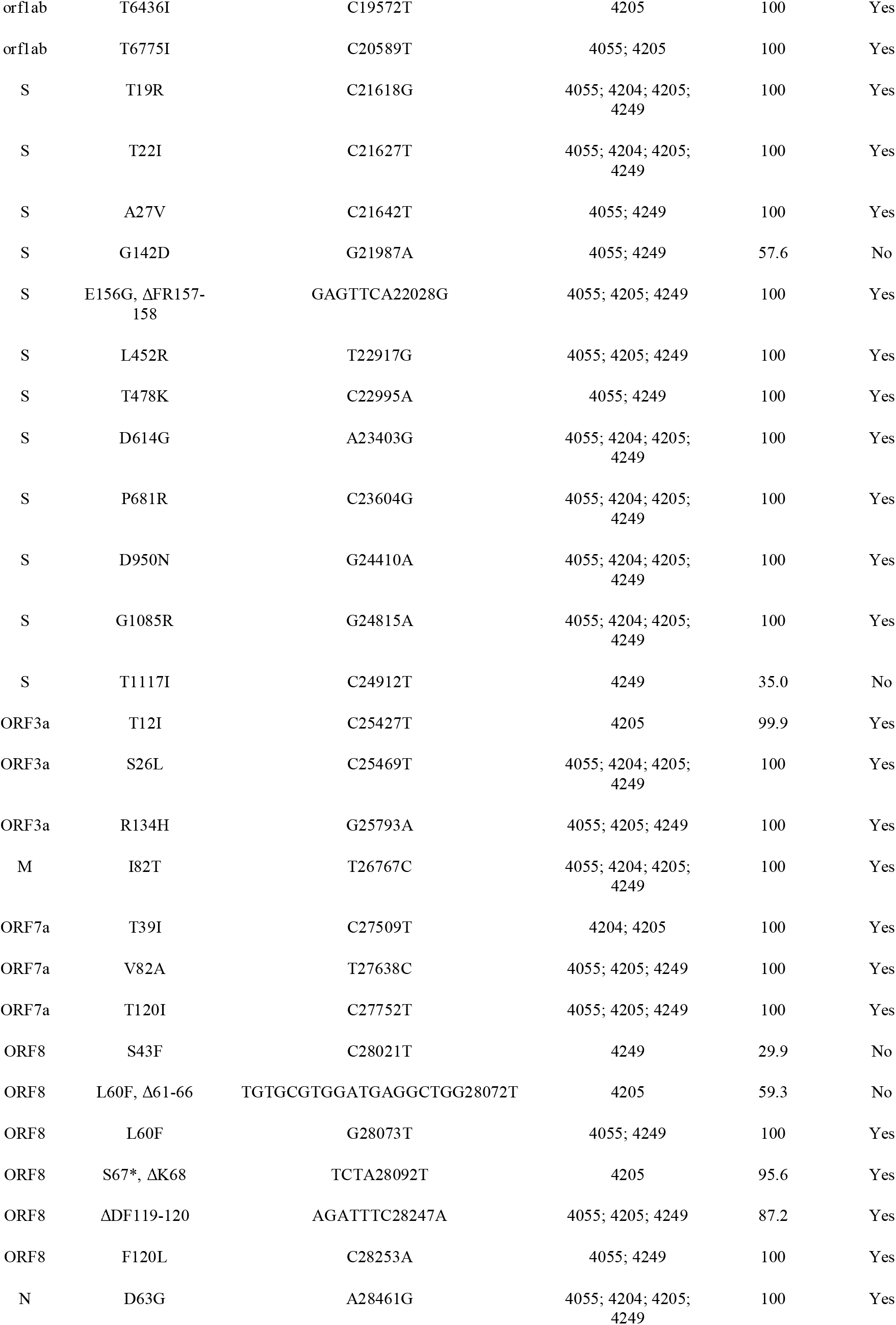
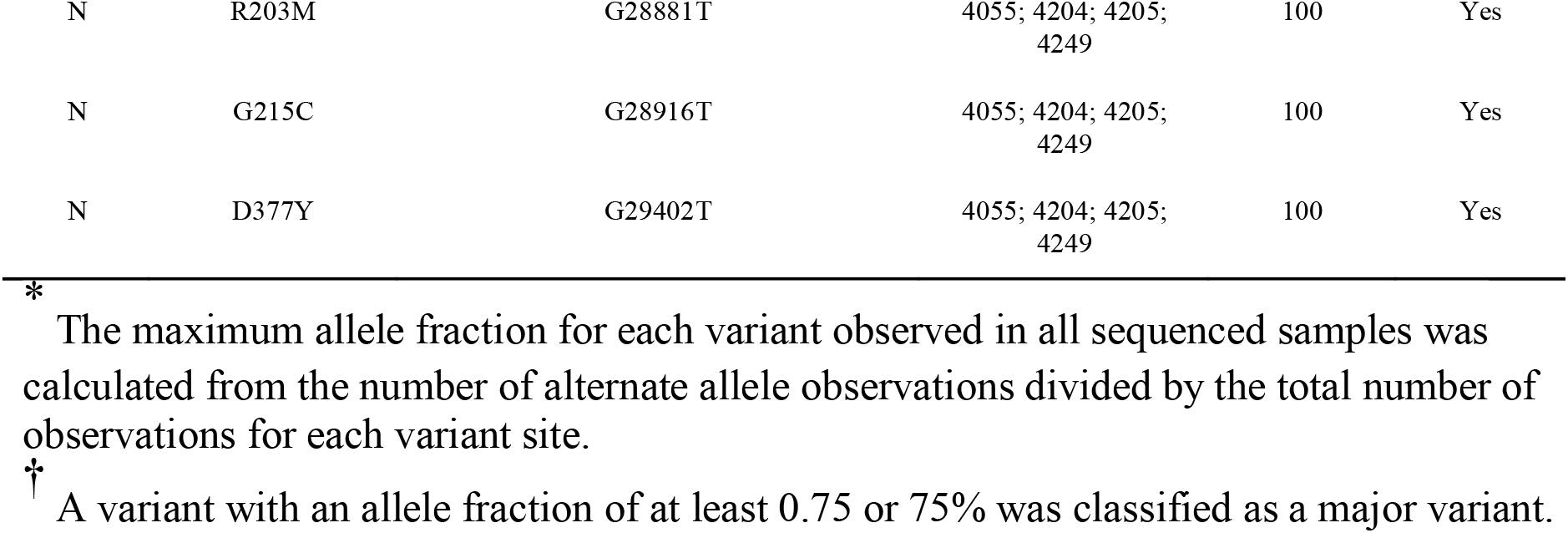
Summary of mutations leading to amino acid changes found in sequences from four SARS-CoV-2 positive white-tailed deer nasal swab samples from southern Québec, Canada, November 6-8 2021

An inframe deletion leading to a stop codon in the ORF8 was observed at S67/K68 (TCTA to T deletion at nucleotide position 28,092) in sample 4205. Additionally, an ORF8 inframe deletion of 6 amino acids at positions 61–66 and L60F mutation were observed in sample 4205 although with an allele fraction of 59% (262/442 observations of TGTGCGTGGATGAGGCTGG to T at nucleotide position 28,072).

As most of the minor alleles were found in sample 4249, it is worth noting that this sample had 15 ambiguous genomic positions with otherwise high completeness (99.1%) and median coverage (1879X). This suggests that more than one SARS-CoV-2 genotype is present within the sequenced sample. This is attributable to either contamination or a mixed infection in the host. Demixing using Freyja v1.3 (https://github.com/andersen-lab/Freyja) revealed the sample was 99.4% AY.44 and 0.1% AY.98. However, no other samples in the same sequencing run contained the 4249-specific variants or were assigned to the AY.98 lineage making contamination less likely. Inter-run contamination is unlikely as a new flow cell was used for each sequencing run and four different, alternating sets of 96 UD Indices were used to barcode the samples in the library pool to help mitigate index hopping. Moreover, all negative controls met quality control parameters, further suggesting contamination is unlikely. As AY.98 is closely related to AY.44 (differing by only 3 ORF1ab amino acid residues) and is inferred as low abundance, this suggests 4249 more likely represents a real mixed infection of two closely related AY.44 genomes.

### Virus isolation

Two of the SARS-CoV-2-positive nasal swabs yielded viable virus when cultured in Vero E6 (**Figure S2**). Resultant sequences from the isolates were comparable to their original nasal swab counterparts; identical mutations found in the original nasal swab sample sequences were also identified in the sequences from the isolates. Differences between sequences from the original samples and the isolated sequences were observed. Changes noted in 4055 after a single passage included S:R683W and an ORF3a deletion V256-259. Five low coverage variants were lost in 4249, which may suggest selection for the more abundant genotype.

### Deer nasal community profile an indicator of COVID-19 infection

The WTD nasal swabs were sequenced by RNAseq, allowing for the taxonomic profiling of the nasal community with Kraken 2 and Bracken (**Table S2**). SARS-CoV-2 was detected in the reads from deer samples that tested positive for SARS-CoV-2 (4055 and 4249) (**Figure S3A**), confirming the RT-PCR results. All swabs were collected and stored in media containing antibiotics and antifungals prior to analysis, which could impact bacterial and fungal profiles. However, the SARS-CoV-2 positive samples cluster together even when excluding SARS-CoV-2 data. Additionally, the RT-PCR-negative deer samples 4192 and 3719 have a divergent microbial profile compared to other negative samples; most notably relative increases in *Cutibacterium acnes* compared to the other deer nasopharyngeal swabs (**Figure S3A**).

### SARS-CoV-2 infection elicits anti-viral, pro- and anti-inflammatory transcriptional response in white-tailed deer

We employed unbiased exploratory transcriptomic analysis to provide insights on WTD host response to SARS-CoV-2 infection. When considering the entire gene expression profile, the deer samples that tested positive for SARS-CoV-2 (4055 and 4249) clustered apart from the negative samples (**Figure S3B**). The negative samples further clustered into two groups, with samples 4192 and 3719 clustering away from the rest (**Figure S3B**). This difference is likely not due to age, sex, or hunting zone since these factors are not unique to the two aforementioned negative samples.

Differential expression analysis identified 316 significant DEGs (adjusted *p*<0.05, >1-fold change), spanning 194 upregulated DEGs and 122 downregulated DEGs (**Figure 3A** and **B** and **Table S3**). Top expressed genes included *LOC110149600* (an inferred ortholog of human *IFITM1*), *LOC110149612* (inferred *IFITM1*), *OAS2, HSH2D*, and *APOBEC3H* (**Table 4** and **Figure 3B**). Notably, several significant DEGs appear to have duplications of human immune genes, including *IFITM1, IFI27L2A, C4a, HNRNPA1*, and *XAF1*, with intriguing implications for the deer immune response. Upon narrowing down to SARS-CoV-2 viral attachment and entry factors that have been identified as important for human COVID-19 infection (e.g., *ACE2* and *TMPRSS2*), only *SIGLEC1* was significantly upregulated in the deer samples (**Figure S4**).

**Table 4.** Top 10 up and down differentially expressed WTD genes ranked by adjusted *p*-value.

| Log<sub>2</sub><br>fold<br>change | Adjusted<br>*p*-value | WTD gene name | Inferred human<br>ortholog* |
| --- | --- | --- | --- |
| Up |  |  |  |
| 5.42 | 1.49x10<sup>-13</sup> | interferon-induced transmembrane protein 1-like (*LOC110149612*) | *IFITM1* |
| 5.50 | 1.84x10<sup>-11</sup> | interferon alpha-inducible protein 27-like protein 2A (*LOC110122860*) |  |
| 4.84 | 2.75x10<sup>-8</sup> | interferon-induced transmembrane protein 1-like (*LOC110149600*) | *IFITM1* |
| 6.34 | 9.71x10<sup>-8</sup> | 2'-5'-oligoadenylate synthetase 2 (*OAS2*) |  |
| 3.96 | 9.12x10<sup>-6</sup> | pleckstrin homology domain-containing family A member 4-like (*LOC110133690*) |  |
| 3.57 | 1.66x10<sup>-5</sup> | hematopoietic SH2 domain containing (*HSH2D*) | *HSH2D* |
| 3.44 | 1.66x10<sup>-5</sup> | DNA dC->dU-editing enzyme APOBEC-3H-like (*LOC110129971*) | *APOBEC3H* |
| 6.12 | 2.70x10<sup>-5</sup> | zonadhesin (*ZAN*) | *ZAN* |
| 4.24 | 6.32x10<sup>-5</sup> | complement C4-A-like (*LOC110139391*) | *C4B_2* |
| 2.95 | 7.22x10<sup>-5</sup> | transporter 1, ATP binding cassette subfamily B member (*TAP1*) | *TAP1* |
| Down |  |  |  |
| -25.49 | 5.00x10<sup>-7</sup> | heterogeneous nuclear ribonucleoprotein A1-like (*LOC110141849*) |  |
| -22.66 | 2.23x10<sup>-5</sup> | cuticlin-2-like (*LOC110149771*) |  |
| -22.56 | 2.31x10<sup>-5</sup> | ornithine decarboxylase antizyme 1-like (*LOC110136472*) |  |
| -21.26 | 9.82x10<sup>-5</sup> | heterogeneous nuclear ribonucleoprotein A1-like (*LOC110143697*) |  |
| -2.90 | 1.42x10<sup>-4</sup> | testis expressed 9 (*TEX9*) | *TEX9* |
| -2.48 | 1.43x10<sup>-4</sup> | G protein subunit alpha 14 (*GNA14*) | *GNA11* |
| -3.82 | 2.68x10<sup>-4</sup> | janus kinase and microtubule interacting protein 2 (*JAKMIP2*) | *JAKMIP2* |
| -2.82 | 5.32x10<sup>-4</sup> | coiled-coil domain containing 181 (*CCDC181*) | *CCDC181* |
| -2.14 | 5.68x10<sup>-4</sup> | homer scaffold protein 2 (*HOMER2*) | *HOMER2* |
| -3.04 | 5.68x10<sup>-4</sup> | histamine receptor H1 (*HRH1*) | *HRH1* |

\*
783 If left blank, no human ortholog could be predicted with OrthoFinder.

**Figure 3.**
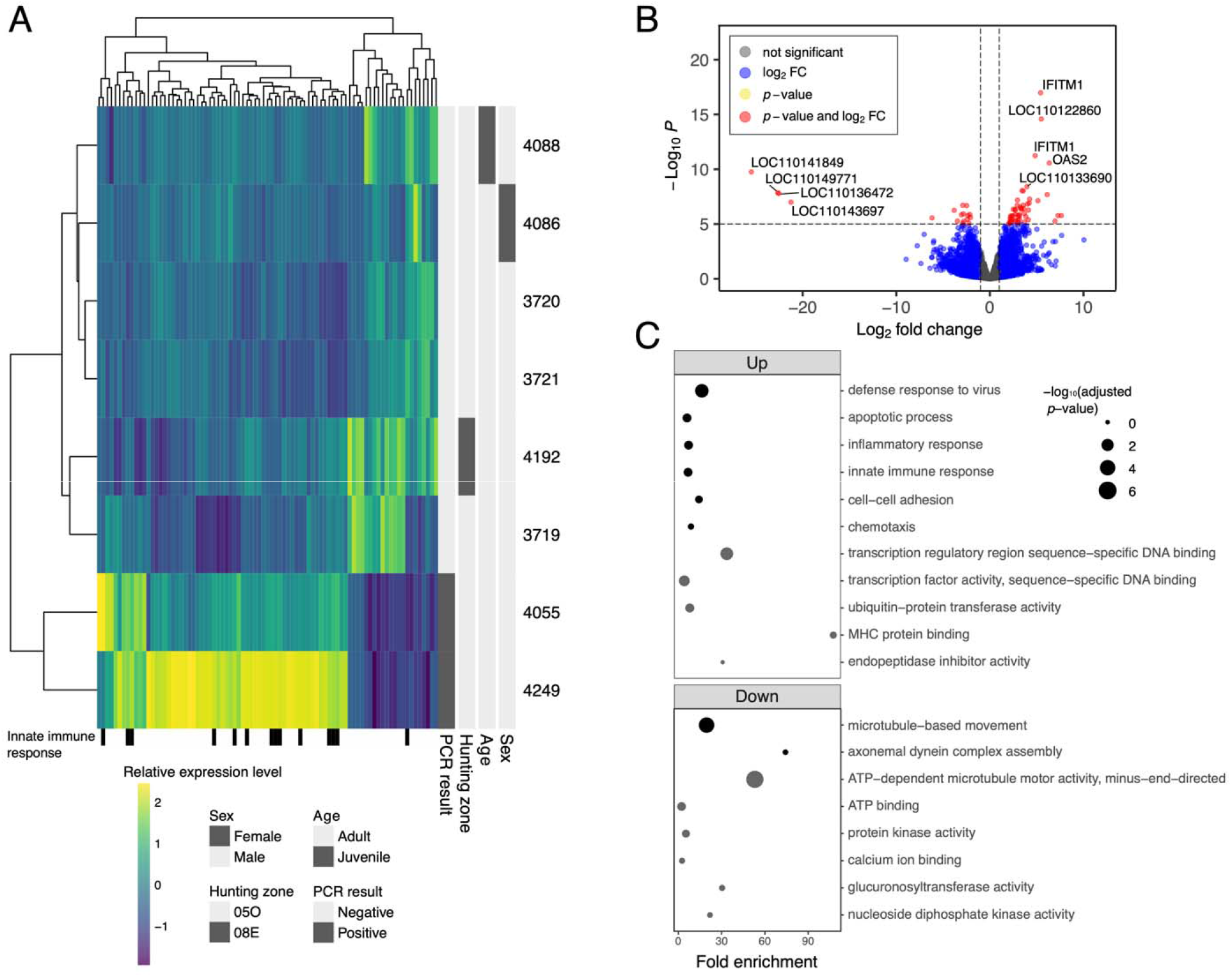
RNAseq analysis of deer nasopharyngeal swabs comparing healthy deer with SARS-CoV-2 infected deer (4055 and 4249). A) Relative expression levels of significant (adjusted *p*-value < 0.05) differentially expressed genes with an absolute fold change greater than 3. Deer genes with human orthologs annotated with the Gene Ontology “innate immune response” term indicated underneath. B) Volcano plot of the DESeq2 differential gene expression analysis results. Where possible, genes were labelled with their inferred human ortholog gene name. The upper *IFITM1* is associated with *LOC110149600* and the lower *IFITM1* is associated with *LOC110149612* in the deer genome. C) Gene Ontology function enrichment of the significant up and down-differentially expressed genes against a deer genome background with DAVID. Only terms from biological processes (black dots) and molecular functions (grey dots) are displayed.

DEG function enrichment analysis using DAVID (**Tables S3-S6**) revealed a change in the expression of key host factors mediating innate immune response (interferon [IFN] signalling, inflammasome, APOBECs), pathology (ciliary function and apoptosis), and permissivity (receptors, attachment, and entry factors) in RT-PCR-positive vs. RT-PCR-negative deer. Significantly enriched Gene Ontology terms among up-regulated DEGs included defence response to virus, apoptotic process, innate immune response, inflammatory response, and chemotaxis, while down-regulated DEGs were involved in microtubule-based movement, axonemal dynein complex assembly, and ATP binding (**Figure 3C**). In humans and mice, the up-regulated DEGs are known to play important roles in type I and type III IFN signalling (e.g., *IRF3, IRF4, IRF5, IRF7, IRF9, INF*_λ_*3, STAT2, IFITM1, IFITM2, IFITM3, IFI6, IFI27L2A*), complement activation (e.g., *C2, C4a*), nucleic acid detection (e.g., *DHX58*), immune cell recruitment (e.g., *CSF1, CCR1, CXCR2, RIPOR2*), apoptosis (e.g., *ELMO, BID, XAF1, DNASE1L3*), and host defence (e.g., *BST2, ZBP1, OAS2, ADAR*). Orthologous DEGs that were downregulated are associated with microtubules (e.g., *DYNC2H1, DYNLRB2, DNAH3/5/7*) that make up extracellular structures such as cilia, likely reflecting host epithelial damage.

### White-tailed deer compared to human SARS-CoV-2 infection response

To identify potential differences in SARS-CoV-2 host responses between humans and deer, we compared the log_2_ fold change of SARS-CoV-2 positive samples to negative samples from human and WTD nasopharyngeal swabs (**Figure 4A** and **Table S8**). The human cohort contained 50 SARS-CoV-2 positive patients (divided into differing levels of disease severity with 18 ICU, 16 non-ICU, and 16 outpatient) and 13 SARS-CoV-2 negative individuals ^53^. When examining all genes associated with innate immune response, there is a weak but statistically significant correlation between deer and human nasopharyngeal log_2_ fold change (*r* = 0.27, *p*-value = 9.49x10^−13^), with a slightly higher correlation observed when comparing against only outpatients (*r* = 0.35, *p*-value = 1.45x10^−20^). For this analysis we only focused on the ∼81% of deer genes with detected human orthologs and did not include deer-specific DEGs. We identified several immune-related genes that were significantly up-regulated in deer and not in the human clinical cohort, including *IFNL1, BST2*, and *IFITM1* (**Figure 4B**). In the data from the Butler et al., (2021) human clinical cohort (RNAseq of naso/oropharyngeal swabs collected from 669 patients), some of these genes are significantly differentially expressed in SARS-CoV-2 positive versus negative patients, but still have low log_2_ fold changes when compared to deer (0.84, 1.93, and 1.94 respectively), although the expression levels of these genes generally increases when comparing patients with higher viral loads than those with none ^60^. Key viral entry and attachment factors for human COVID-19 infection were also examined in this way (**Figure S5**), with five of the set (*ACE2, BSG, CTSV, MMP2*, and *SIGLEC1*) being significant DEGs in the human nasopharyngeal samples compared to only one (*SIGLEC1*) in the deer samples.

**Figure 4.**
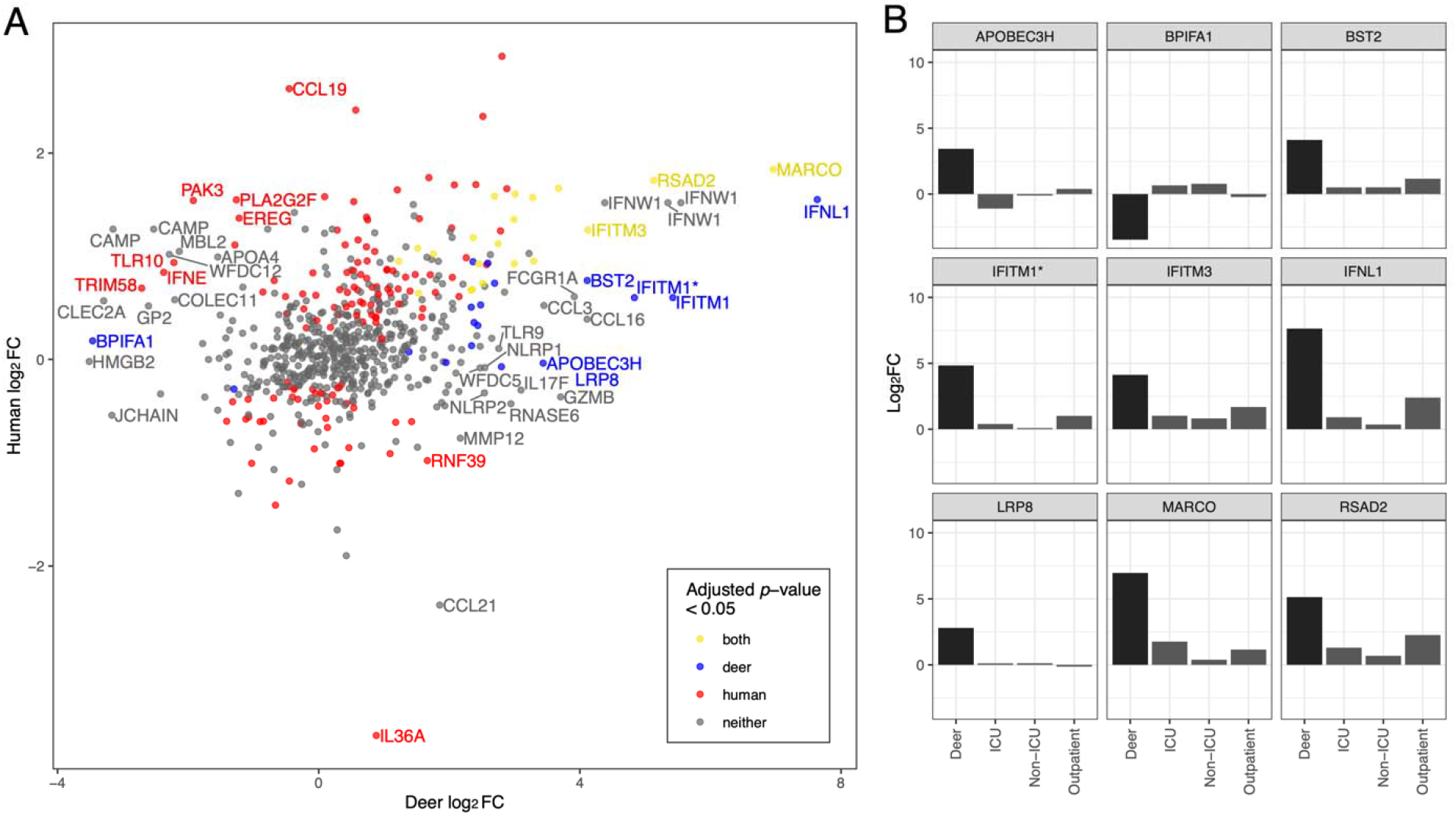
Comparison of the WTD versus human host response to SARS-CoV-2 infection. A) The WTD expression fold change is plotted here against the human expression fold change from comparable human samples based on the results of an orthology analysis between WTD and human. The genes are labelled with the respective human gene name, if the absolute difference in log_2_ fold change values between the human and deer expression profiles is greater than 2.5. Due to duplicated genes in the deer genome, some points are labelled with the same human gene name. From left to right, *CAMP* genes are associated with *LOC110152352, LOC110152353*, and *LOC110152344* in the deer genome, *IFNW1* genes are associated with *LOC110144825* and *LOC110144833* and *IFITM1* genes are associated with *LOC110149600* and *LOC110149612*. Points are coloured based on significance in their respective differentially gene expression analysis. Genes with very low expression values are not given a *p*-value in the DESeq2 analysis and are not considered significant. B) Genes that are significantly differentially expressed in the deer SARS-CoV-2 infection and have an absolute difference in log_2_ fold change values between the human and deer infections greater than 2.5 are plotted here with their corresponding log_2_ fold change values across different levels of human disease severity. ICU, non-ICU, and outpatient are comparisons of human nasopharyngeal swabs between patients from these respective settings, and healthy patients. Genes are labelled with their human ortholog gene name. *IFITM1* here corresponds to *LOC110149600* in the deer genome.

**Figure 5.**
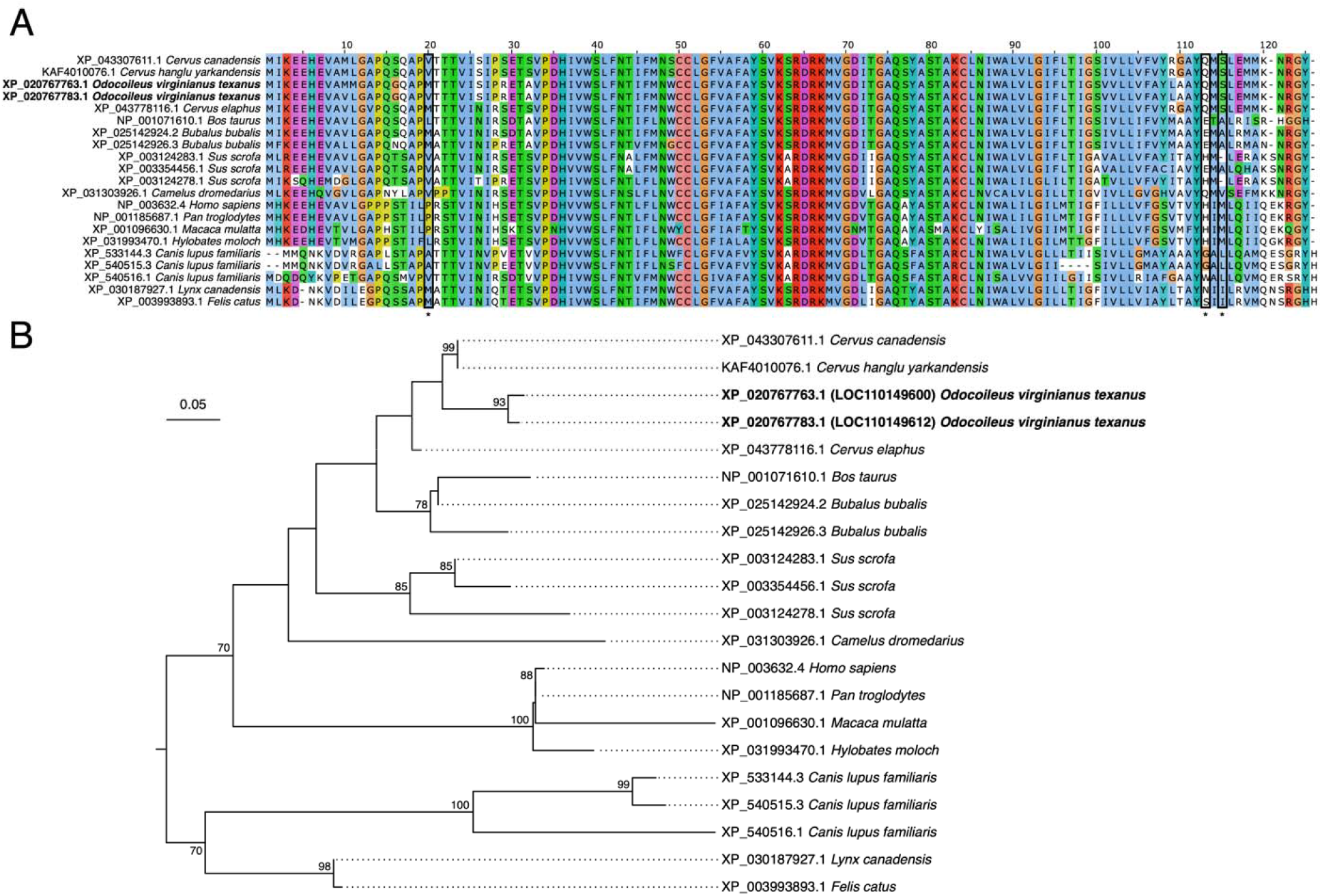
Phylogenetic analysis of IFITM1 in mammals reveals lineage-specific duplications in deer and evolutionary diversification. A) Alignment of IFITM1 from select boreoeutherian mammals. An extended N-terminal start to XP_043307611.1 (*Cervus canadensis*) and XP_025142924.2 (*Bubalus bubalis*) has been trimmed from the alignment. Boxed and asterisked positions 20, 113 and 115 were identified as sites of diversifying selection. Alignment is coloured with the Clustal colour scheme with poorly conserved residues left white using Jalview v2.11.2.6. B) A mid-point rooted tree of IFITM1 from select boreoeutherian mammals. Bootstrap values less than 70 were not displayed.

Orthology analysis between human and the *texanus* subspecies of WTD revealed two interferon induced transmembrane 1 (*IFITM1*) genes in WTD, compared to a single copy in humans. Both of these genes were significantly over-expressed in WTD with SARS-CoV-2 infections (log_2_ fold change of 4.84 and 5.42 with adjusted *p*-values of 2.82x10^−8^ and 1.53x10^−13^ for *LOC110149600* and *LOC110149612*, respectively). However, in the human cohort, *IFITM1* was not significantly differentially expressed, and log_2_ fold change values were low. Other WTD genomes (strains 20LAN1187 and brownington 1) were also confirmed to have two *IFITM1* genes, with 99% similarity between the sequenced *LOC110149600* genes (2-4 nucleotide differences) and 99% similarity between the *LOC110149612* genes (6-12 nucleotide differences). *LOC110149600* and *LOC110149612* are separated on the genome, with an average distance of 26,485 +/-486 base pairs apart. The *LOC110149612* transcript has an extended N-terminus compared to *LOC110149600* but with respect to their protein sequences, there are only three amino acid substitutions between them: valine/methionine (position 10), leucine/methionine (position 11), and valine/methionine (position 20) for *LOC110149612* and *LOC110149600*, respectively. Near identical IFITM1 sequences are also found in the other two WTD genomes. The identity between the human and WTD IFITM1 proteins is much lower at 66-68%, with particular differences in the N-terminus including several unique proline substitutions that could impact protein structure. A Fixed Effects Likelihood analysis identified positions 20, 113, and 115 to have evidence of diversifying selection (**Table S9**), with all three of these positions differing between humans and WTD. A phylogenetic tree based on IFITM1 proteins from boreoeutherian mammals shows multiple instances of *IFITM1* gene duplication, all showing the extensive diversification *IFITM1* has undergone throughout mammals and the substantial differences between human and WTD IFITM1.

## Discussion

In this report, we detected SARS-CoV-2 in 1.6% (95% CI 0.5–4.2%) of nasal swabs from sampled deer in the Estrie region of southern Québec; viral sequences were assigned to lineage AY.44, a sublineage of the Delta variant of concern (VOC). Delta was the predominant circulating VOC at the time these animals were sampled, which is suggestive of a more recent spillover event. However, several unique mutations observed in the WTD sequences were not observed in the closely related AY.44 sequences from GISAID, supporting sustained deer-deer transmission. The WTD derived sequences from this study were most closely related to SARS-CoV-2 sequences from humans in neighbouring Vermont, USA and one sequence from a human in Québec (**Figure 2**). Although we included SARS-CoV-2 sequences from humans from Québec in the analysis, it is unknown if any of these sequences were from the Estrie region. While the Estrie region borders Vermont, to our knowledge, there is no evidence of SARS-CoV-2 in WTD that has been reported in Vermont to date ^61^.

Whole SARS-CoV-2 genome sequences were found to contain two S gene mutations (S:T22I and S:A27V) that were different between deer SARS-CoV-2 and the most closely related AY.44 sequences from GISAID. These changes are both located in the N terminal domain (NTD) of S1, which harbours antigenic and glycan-binding sites, and may interact with auxiliary receptors ^62^. Changes at amino acid position 22 and 27 have been noted early in the pandemic, with up to four different amino acid variants at position 27; this plasticity is suggestive of an adaptive evolutionary role ^63,64^. We have since detected the S:T22I mutation in a highly divergent, deer-adapted SARS-CoV-2 from Ontario ^10^, further supporting a relevant role for changes in this region of the S protein. Additionally, in one deer (4205), two inframe deletions in ORF8 were observed. The ORF8 gene is hypervariable and encodes for a non-essential accessory protein and has been shown to downregulate the major histocompatibility complex class I through autophagic degradation, thus impairing cytotoxic T cell responses during SARS-CoV-2, but not SARS-CoV infection ^65^. It may also contribute to immune evasion through interferon antagonism ^66,67^. More recently, ORF8 protein was found to purportedly mimic host interleukin-17 that contributes to severe inflammation in COVID-19 ^68^. Truncation of ORF8 arising from nonsense mutations and deletions have been previously observed in both human and animal derived sarbecoviruses (SARS-CoV and SARS-CoV-2) ^66,69–71^ and may be associated with milder disease through enhanced T cell functions. This underscores the potential role of ORF8 in SARS-CoV-2 adaptation.

All RT-PCR-positive WTD were identified in the Estrie region (**Table 1; Figure 1**). This is unsurprising given the reported spatial clustering of SARS-CoV-2 in WTD in other regions ^1,5,6^. While it is presently unclear how the deer acquired SARS-CoV-2 infection, there are several notable differences between the sampled regions that may contribute to the spatial heterogeneity of infection: 1) WTD population density and hunter harvest are greater in Estrie (13-15 deer/km^2^) compared to the Laurentides (∼1 deer/km^2^) (MELCCFP, unpublished data), 2) the sampled regions in Estrie have higher human population density compared to sampled regions in the Laurentides (**Figure 1A**), and 3) COVID-19 positivity in humans was greater in Estrie (4.2%) during the study period compared to the Laurentides (2.3%) ^72^. More longitudinal surveillance of SARS-CoV-2 in WTD is needed to understand the epidemiology of the virus in this species and how it relates to WTD ecology and transmission dynamics in sympatric humans. These are important considerations for understanding the potential role of WTD as a maintenance population or reservoir for SARS-CoV-2.

Based on previous experimental work, productive viral replication is limited to the upper respiratory tract with shedding of infectious virus in nasal secretions of infected WTD ^3,9^. We successfully isolated viable SARS-CoV-2 from two RT-PCR-positive nasal swabs indicating infectivity. This suggests there is a potential risk of contact with infectious SARS-CoV-2 from deer, including when handling and processing WTD carcasses ^1^. Notably, there is only one report of an isolated, unsustained deer-to-human transmission event to date ^10^. These findings warrant increased awareness of the risks associated with human contact with free-living and captive WTD ^73^.

Previous work characterizing the WTD host response to SARS-CoV-2 infection is limited. Davila and colleagues conducted a comparative transcriptomics analysis of SARS-CoV-2 infected human and deer primary respiratory epithelial cells from the trachea, finding evidence of divergent early innate immune response ^74^. The present study is the first to explore the host response in WTD naturally infected with SARS-CoV-2. Host expression patterns in SARS-CoV-2 infected WTD were associated with the innate immune response, including signalling pathways related to anti-viral and pro-inflammatory signalling and host damage. There was evidence for type I and III IFN responses (e.g., *IRF3, IRF4, IRF5, IRF7, IRF9, INF*_λ_*3, STAT2, IFITM1, IFITM2, IFITM3, IFI6, IFI27L2A*). The type I IFN response is an important aspect of rapidly controlling viral infection, but dysregulation is associated with severe illness and pathology. Type III IFNs are considered to produce a more localized response to infection in comparison to the systemic inflammatory response often induced by type I IFN; type III IFNs result in prolonged expression of ISGs ^75^. Upregulation of pro-inflammatory antagonists were also observed. For example, there was increased expression of two NF-KB inhibitors (e.g. *NFKBID, NFKBIZ*) which could indicate a mechanism to reduce deleterious inflammation or indicate a switch to resolution of inflammation phase (e.g., tissue repair) ^76^. Additionally, several genes involved in antiviral response and inflammation homeostasis were found to have multiple DEG copies (e.g., *IFITM1, XAF1*) which may also contribute to protecting the host from inflammation ^77,78^.

When examining all genes associated with innate immune response, there is a weak but statistically significant correlation between deer and human log_2_ fold change, with the strongest correlation observed between deer and outpatients. Previous studies have reported that WTD infected with SARS-CoV-2 do not present with overt signs of infection or pathology; only minor pathological changes associated with rhinitis and marked attenuation of the respiratory epithelium have been observed in experimentally infected WTD ^3,7,9^. However, duration of infection for each individual host is unknown, limiting the inferences that can be made regarding the dynamics of innate immune responses. Additionally, it is unclear whether infection with SARS-CoV-2 results in any sublethal effects (e.g., condition, winter survival, reproduction) despite no overt signs of infection.

There were also DEGs with substantial differences within the innate immune response gene set, notably *MARCO, IFNL1, IFITM1, IFITM3, RSAD2, BST2, APOBEC3H, LRP8*, and *BPIFA1*. These genes had a large difference (>2.5) in log_2_ fold change values between the human and deer DEGs while also being significantly differentially expressed in deer. These data indicate that the deer nasal epithelium expressed genes that encode antiviral proteins, including the *IFITM* family, in response to SARS-CoV-2 infection. IFITMs are membrane proteins that restrict viral entry for a broad spectrum of enveloped viruses ^79,80^. In contrast, SARS-CoV-2 has been shown to hijack IFITMs for efficient cellular entry, a process mediated by specific interactions between the N-terminal region of IFITMs and the viral S protein ^78,80^. *IFITM1* is an intriguing case as it has undergone a lineage-specific duplication resulting in two copies in deer, both of which were significantly upregulated in this study. The amino acid changes between the two WTD IFITM1 are present in other mammalian (e.g. other deer in the *Cervus* genus and Felinae) lineages and are thus not unique to WTD deer, but do appear to have evolved independently on several occasions. More importantly, the N- and C-terminal ends of both IFITM1 possess unique differences in deer compared to human IFITM1. We speculate that these differences might modulate differences in antiviral properties or efficiency of IFITM1 in deer (and other animals) versus humans, as previously seen in the IFITM3 family in primates ^81^, and are an important future target for characterization of the SARS-CoV-2 infection response in WTD.

There are several limitations for this study that should be considered. First, while leveraging the regular WTD hunting season resulted in a large number of samples, the present work was conducted over a short time period and a relatively small geographic region. Therefore, our study represents a snapshot in time and space. Future work should aim to obtain longitudinal data to investigate maintenance of SARS-CoV-2 in Québec WTD populations and assess for spatiotemporal patterns in pathogen ecology. Second, samples analyzed in this study were derived from harvested deer and were therefore collected post-mortem. Although 98% of samples were collected within 48 hours of harvest, it is possible that inhibitors or sample degradation occurred between harvest and sample collection. Lastly, our study only focuses on free-ranging WTD populations and we therefore cannot make inferences about SARS-CoV-2 in captive conspecifics.

Surveillance for SARS-CoV-2 in wildlife is ongoing across Canada. Our findings underscore that longitudinal surveillance efforts in WTD in Québec and across Canada are warranted. We provide preliminary insights into unique transcriptional responses in white-tailed deer with SARS-CoV-2 infection. Further work is needed to understand how the virus is transmitted from humans to deer, how efficiently and sustainably the virus is transmitted among deer in a natural setting, and how viral adaptations occur in WTD. Additionally, more longitudinal epidemiological and ecological data is needed to better understand whether WTD truly represent a competent maintenance population or reservoir for SARS-CoV-2. Ongoing coordinated and cross-disciplinary efforts are required to ensure a One Health approach is applied to this critical pandemic challenge by informing evidence-based decision-making for human and animal health.

## Acknowledgements

We thank the hunters for the deer submissions. Many thanks to the technicians and biologists who assisted with sample collection (Matthew Rokas-Bérubé, Yannicia Fréchette-Hudon, Catherine Greaves, Charles-Étienne Gagnon, Marylou Meyer, Camille Klein, Stéphane Lamoureux, Yannick Bilodeau, François Lebel, Sophie Plante, and Isabelle Laurion), and the technicians that assisted with sample analysis (Emily Chien, Mathieu Pinette, Winfield Yim, Melissa Goolia, and Nikki Toledo). We are grateful for the work of D. Bulir in developing the UTR gene target used in the Sunnybrook Research Institute RT-PCR analysis. This work was supported by the Public Health Agency of Canada, and Canadian Institutes of Health Research Operating Grant: Emerging COVID-19 Research Gaps and Priorities (466984). J.D.K. is supported by an AMMI Canada/BioMérieux Fellowship in Microbial Diagnostics. Funding and computing resources for F.M were provided by the Shared Hospital Laboratory, Dalhousie University, and the Donald Hill Family. O.L. was supported by the Canadian Safety and Security Program, Laboratories Canada, and Canadian Food Inspection Agency (CFIA) Genomics Development and Research Initiative. A.B. also acknowledges support from Natural Sciences and Engineering Research Council of Canada (NSERC), Canadian Institutes of Health Research (CIHR), and Coronavirus Variants Rapid Response Network (CoVaRR-Net). VIDO receives operational funding for its CL3 facility (InterVac) from the Canada Foundation for Innovation through the Major Science Initiatives. VIDO also receives operational funding from the Government of Saskatchewan through Innovation Saskatchewan and the Ministry of Agriculture.

## Author contributions

Conceptualization: J.D.K., A.M., M.G., J.B., T.B., B.P., S.M.

Sample collection: J.D.K., A.M., M.G.

Laboratory analysis: J.D.K, P.A., J.B.S., H.Y.C., K.N., L.Y., L.R.L., B.P.

Data analysis/Investigation: J.D.K., B.L., P.K., F.M., O.L.

Writing – Original Draft: J.D.K., B.L.

Writing – Review & Editing: All authors

Visualization: J.D.K., B.L., P.K., A.C.D., O.L.

Supervision: L.R.L., A.C.D., O.L., B.P., S.M.

Funding acquisition: A.M., M.G., S.M.

## Data availability

Sequence data from the three SARS-CoV-2 viruses from white-tailed deer sequenced in this study are deposited in GISAID (https://www.gisaid.org/) under Accession numbers EPI_ISL_10169675 (hCoV-19/Canada/QC-WTD-qxic4205/2021), EPI_ISL_10170149 (hCoV-19/Canada/QC-WTD-qxic4249/2021), and EPI_ISL_10168587 (hCoV-19/Canada/QC-WTD-qxic4055/2021).

## References

1. Caserta, L. C. et al. White-tailed deer (Odocoileus virginianus) may serve as a wildlife reservoir for nearly extinct SARS-CoV-2 variants of concern. Proc. Natl. Acad. Sci. 120, e2215067120 (2023).

2. Chandler, J. C. et al. SARS-CoV-2 exposure in wild white-tailed deer (Odocoileus virginianus). Proc. Natl. Acad. Sci. 118, (2021).

3. Cool, K. et al. Infection and transmission of ancestral SARS-CoV-2 and its alpha variant in pregnant white-tailed deer. Emerg. Microbes Infect. 0, 1–39 (2021).

4. Hale, V. L. et al. SARS-CoV-2 infection in free-ranging white-tailed deer. Nature 602, 481–486 (2022).

5. Kuchipudi, S. V. et al. Multiple spillovers from humans and onward transmission of SARS-CoV-2 in white-tailed deer. Proc. Natl. Acad. Sci. 119, (2022).

6. Marques, A. D. et al. Multiple Introductions of SARS-CoV-2 Alpha and Delta Variants into White-Tailed Deer in Pennsylvania. mBio 13, e02101–22 (2022).

7. Martins, M. et al. From Deer-to-deer: SARS-CoV-2 is efficiently transmitted and presents broad tissue tropism and replication sites in white-tailed deer. PLOS Pathog. 18, e1010197 (2022).

8. Palermo, P. M., Orbegozo, J., Watts, D. M. & Morrill, J. C. SARS-CoV-2 neutralizing antibodies in white-tailed deer from Texas. Vector-Borne Zoonotic Dis. 22, 62–64 (2022).

9. Palmer, M. V. et al. Susceptibility of white-tailed deer (Odocoileus virginianus) to SARS-CoV-2. J. Virol. 95, (021).

10. Pickering, B. et al. Divergent SARS-CoV-2 variant emerges in white-tailed deer with deer-to-human transmission. Nat. Microbiol. 7, 2011–2024 (2022).

11. Roundy, C. M. et al. High seroprevalence of SARS-CoV-2 in white-tailed deer (Odocoileus virginianus) at one of three captive cervid facilities in Texas. Microbiol. Spectr. 10, e00576–22 (2022).

12. Tan, C. C. S. et al. Transmission of SARS-CoV-2 from humans to animals and potential host adaptation. Nat. Commun. 13, 2988 (2022).

13. Bashor, L. et al. SARS-CoV-2 evolution in animals suggests mechanisms for rapid variant selection. Proc. Natl. Acad. Sci. 118, e2105253118 (2021).

14. Larsen, H. D. et al. Preliminary report of an outbreak of SARS-CoV-2 in mink and mink farmers associated with community spread, Denmark, June to November 2020. Eurosurveillance 26, 2100009 (2021).

15. Wei, C. et al. Evidence for a mouse origin of the SARS-CoV-2 Omicron variant. J. Genet. Genomics 48, 1111–1121 (2021).

16. Banerjee, A. et al. Novel insights into immune systems of bats. Front. Immunol. 11, 26 (2020).

17. Hiller, M. et al. Reference-quality bat genomes illuminate adaptations to viral tolerance and disease resistance. Preprint at Research Square https://doi:10.21203/rs.3.rs-2557682/v1 (2023).

18. Irving, A. T., Ahn, M., Goh, G., Anderson, D. E. & Wang, L.-F. Lessons from the host defences of bats, a unique viral reservoir. Nature 589, 363–370 (2021).

19. Subudhi, S., Rapin, N. & Misra, V. Immune system modulation and viral persistence in bats: understanding viral spillover. Viruses 11, 192 (2019).

20. Becker, D. J. & Banerjee, A. Coupling field and laboratory studies of immunity and infection in zoonotic hosts. Lancet Microbe 4, E285–E287 (2023).

21. Mandl, J. N. et al. Reservoir host immune responses to emerging zoonotic viruses. Cell 160, 20–35 (2015).

22. Gagnier, M., Laurion, I. & DeNicola, A. J. Control and surveillance operations to prevent chronic wasting disease establishment in free-ranging white-tailed deer in Québec, Canada. Animals 10, 283 (2020).

23. LeBlanc, J. J. et al. Real-time PCR-based SARS-CoV-2 detection in Canadian laboratories. J. Clin. Virol. 128, 104433 (2020).

24. Lu, X. et al. US CDC Real-Time Reverse Transcription PCR panel for detection of severe acute respiratory syndrome coronavirus 2. Emerg. Infect. Dis. 26, 1654–1665 (2020).

25. World Organization for Animal Health. Considerations for sampling, testing, and reporting of SARS-CoV-2 in animals. [cited 2023 May 5]. https://www.oie.int/app/uploads/2021/03/a-sampling-testing-and-reporting-of-sars-cov-2-in-animals-3-july-2020.pdf. (2020).

26. Agresti, A. & Coull, B. A. Approximate is better than “exact” for interval estimation of binomial proportions. Am. Stat. 52, 119–126 (1998).

27. Statistics Canada. hPopulation and dwelling count highlight tables, 2016 Census [cited 2021 Dec 07]. https://www12.statcan.gc.ca/census-recensement/2016/dp-pd/hlt-fst/pd-pl/Table.cfm?Lang=Eng&T=101&S=50&O=A#2016A000224

28. Ministère des Forêts, de la Faune et des Parcs (MFFP). Season 2020 - White-tailed deer harvest per hunting zone. 4 (2020).

29. Di Tommaso, P. et al. Nextflow enables reproducible computational workflows. Nat. Biotechnol. 35, 316–319 (2017).

30. Ewels, P. A. et al. The nf-core framework for community-curated bioinformatics pipelines. Nat. Biotechnol. 38, 276–278 (2020).

31. Patel, H. et al. nf-core/viralrecon: nf-core/viralrecon v2.5 - Manganese Monkey. Zenodo https://doi:10.5281/ZENODO.3901628 (2022)

32. Elbe, S. & Buckland-Merrett, G. Data, disease and diplomacy: GISAID’s innovative contribution to global health. Glob. Chall. 1, 33–46 (2017).

33. Khare, S. et al. GISAID’s role in pandemic response. China CDC Wkly. 3, 1049–1051 (2021).

34. Shu, Y. & McCauley, J. GISAID: Global initiative on sharing all influenza data –from vision to reality. Eurosurveillance 22, 30494 (2017).

35. Turakhia, Y. et al. Ultrafast Sample placement on Existing tRees (UShER) enables real-time phylogenetics for the SARS-CoV-2 pandemic. Nat. Genet. 53, 809–816 (2021).

36. Aksamentov, I., Roemer, C., Hodcroft, E. & Neher, R. Nextclade: clade assignment, mutation calling and quality control for viral genomes. J. Open Source Softw. 6, 3773 (2021).

37. Nguyen, L.-T., Schmidt, H. A., von Haeseler, A. & Minh, B. Q. IQ-TREE: A fast and effective stochastic algorithm for estimating maximum-likelihood phylogenies. Mol. Biol. Evol. 32, 268–274 (2015).

38. Minh, B. Q. et al. IQ-TREE 2: New models and efficient methods for phylogenetic inference in the genomic era. Mol. Biol. Evol. 37, 1530–1534 (2020).

39. Kalyaanamoorthy, S., Minh, B. Q., Wong, T. K. F., von Haeseler, A. & Jermiin, L. S. ModelFinder: fast model selection for accurate phylogenetic estimates. Nat. Methods 14, 587–589 (2017).

40. Cock, P. J. A. et al. Biopython: freely available Python tools for computational molecular biology and bioinformatics. Bioinforma. Oxf. Engl. 25, 1422–1423 (2009).

41. Yu, G., Smith, D. K., Zhu, H., Guan, Y. & Lam, T. T.-Y. ggtree: an r package for visualization and annotation of phylogenetic trees with their covariates and other associated data. Methods Ecol. Evol. 8, 28–36 (2017).

42. Nasir, J. A. et al. A comparison of whole genome sequencing of SARS-CoV-2 using amplicon-based sequencing, random hexamers, and bait capture. Viruses 12, 895 (2020).

43. Chen, S., Zhou, Y., Chen, Y. & Gu, J. fastp: an ultra-fast all-in-one FASTQ preprocessor. Bioinforma. Oxf. Engl. 34, i884–i890 (2018).

44. Dobin, A. et al. STAR: ultrafast universal RNA-seq aligner. Bioinforma. Oxf. Engl. 29, 15–21 (2013).

45. O’Hara, E. et al. Neural transcriptomic signature of chronic wasting disease in white-tailed deer. BMC Genomics 23, 69 (2022).

46. Bushnell, B. BBMap: A Fast, Accurate, Splice-Aware Aligner. https://www.osti.gov/biblio/1241166 (2014).

47. Wood, D. E., Lu, J. & Langmead, B. Improved metagenomic analysis with Kraken 2. Genome Biol. 20, 257 (2019).

48. Lu, J., Breitwieser, F. P., Thielen, P. & Salzberg, S. L. Bracken: estimating species abundance in metagenomics data. PeerJ Comput. Sci. 3, e104 (2017).

49. Patro, R., Duggal, G., Love, M. I., Irizarry, R. A. & Kingsford, C. Salmon provides fast and bias-aware quantification of transcript expression. Nat. Methods 14, 417–419 (2017).

50. Love, M. I., Huber, W. & Anders, S. Moderated estimation of fold change and dispersion for RNA-seq data with DESeq2. Genome Biol. 15, 550 (2014).

51. Sherman, B. T. et al. DAVID: a web server for functional enrichment analysis and functional annotation of gene lists (2021 update). Nucleic Acids Res. 50, W216–221 (2022).

52. Emms, D. M. & Kelly, S. OrthoFinder: phylogenetic orthology inference for comparative genomics. Genome Biol. 20, 238 (2019).

53. Luc, J. et al. Comparative transcriptomic analysis of nasopharyngeal swabs from individuals with and without COVID-19. Figshare https://doi:10.6084/M9.FIGSHARE.22704403.V1. (2023)

54. Liberzon, A. et al. The Molecular Signatures Database (MSigDB) hallmark gene set collection. Cell Syst. 1, 417–425 (2015).

55. Edgar, R. C. MUSCLE: multiple sequence alignment with high accuracy and high throughput. Nucleic Acids Res. 32, 1792–1797 (2004).

56. Stamatakis, A. RAxML version 8: a tool for phylogenetic analysis and post-analysis of large phylogenies. Bioinformatics 30, 1312–1313 (2014).

57. Kosakovsky Pond, S. L. & Frost, S. D. W. Not so different after all: a comparison of methods for detecting amino acid sites under selection. Mol. Biol. Evol. 22, 1208–1222 (2005).

58. Weaver, S. et al. Datamonkey 2.0: A modern web application for characterizing selective and other evolutionary processes. Mol. Biol. Evol. 35, 773–777 (2018).

59. World Organization for Animal Health. SARS-CoV-2 in animals –situation report 8. [cited 2023 May 5]. https://www.oie.int/app/uploads/2022/01/sars-cov-2-situation-report-8.pdf. (2021).

60. Butler, D. et al. Shotgun transcriptome, spatial omics, and isothermal profiling of SARS-CoV-2 infection reveals unique host responses, viral diversification, and drug interactions. Nat. Commun. 12, 1660 (2021).

61. Despres, H. W. et al. Surveillance of Vermont wildlife in 2021-2022 reveals no detected SARS-CoV-2 viral RNA. 2023.04.25.538264 Preprint at bioRxiv https://doi.org/10.1101/2023.04.25.538264 (2023).

62. Harvey, W. T. et al. SARS-CoV-2 variants, spike mutations and immune escape. Nat. Rev. Microbiol. 19, 409–424 (2021).

63. Long, S. W. et al. Molecular Architecture of Early Dissemination and Massive Second Wave of the SARS-CoV-2 Virus in a Major Metropolitan Area. mBio 11, e02707–20 (2020).

64. Saputri, D. S. et al. Flexible, Functional, and Familiar: Characteristics of SARS-CoV-2 Spike Protein Evolution. Front. Microbiol. 11, (2020).

65. Zhang, Y. et al. The ORF8 protein of SARS-CoV-2 mediates immune evasion through down-regulating MHC-?. Proc. Natl. Acad. Sci. 118, e2024202118 (2021).

66. Li, J.-Y. et al. The ORF6, ORF8 and nucleocapsid proteins of SARS-CoV-2 inhibit type I interferon signaling pathway. Virus Res. 286, 198074 (2020).

67. Moriyama, M., Lucas, C., Monteiro, V. S. & Iwasaki, A. SARS-CoV-2 variants do not evolve to promote further escape from MHC-I recognition. Preprint at bioRxiv http://biorxiv.org/lookup/doi/10.1101/2022.05.04.490614 (2022).

68. Wu, X. et al. Viral mimicry of interleukin-17A by SARS-CoV-2 ORF8. mBio 13, e00402–22 (2022).

69. Fong, S.-W. et al. Robust Virus-specific adaptive immunity in COVID-19 patients with SARS-CoV-2 Δ382 variant infection. J. Clin. Immunol. 42, 214–229 (2022).

70. Su, Y. C. F. et al. Discovery and genomic characterization of a 382-nucleotide deletion in ORF7b and ORF8 during the early evolution of SARS-CoV-2. mBio 11, e01610–20 (2020).

71. Young, B. E. et al. Effects of a major deletion in the SARS-CoV-2 genome on the severity of infection and the inflammatory response: an observational cohort study. The Lancet 396, 603–611 (2020).

72. Institut national de & santé publique du Québec (INSPQ). Données COVID-19 par région sociosanitaire [cited 2023 May 5]. https://www.inspq.qc.ca/covid-19/donnees/par-region

73. Public Health Agency of Canada. Animals and COVID-19. [cited 2023 May 5]. https://www.canada.ca/en/public-health/services/diseases/2019-novel-coronavirus-infection/prevention-risks/animals-covid-19.html

74. Davila, K. M. S. et al. How do deer respiratory epithelial cells weather the initial storm of SARS-CoV-2? Preprint at bioRxiv https://doi.org/10.1101/2023.04.24.538130 (2023).

75. Broadbent, L. et al. An endogenously activated antiviral state restricts SARS-CoV-2 infection in differentiated primary airway epithelial cells. PLOS ONE 17, e0266412 (2022).

76. Lieberman, N. A. P. et al. In vivo antiviral host transcriptional response to SARS-CoV-2 by viral load, sex, and age. PLOS Biol. 18, e3000849 (2020).

77. Han, Y. et al. XAF1 Protects host against emerging RNA viruses by stabilizing IRF1-dependent antiviral immunity. J. Virol. 96, e0077422 (2022).

78. Shi, G. et al. Opposing activities of IFITM proteins in SARS□CoV□2 infection. EMBO J. 40, e106501 (2021).

79. Buchrieser, J. et al. Syncytia formation by SARS-CoV-2-infected cells. EMBO J. 39, e106267 (2020).

80. Prelli Bozzo, C. et al. IFITM proteins promote SARS-CoV-2 infection and are targets for virus inhibition in vitro. Nat. Commun. 12, 4584 (2021).

81. Compton, A. A. et al. Natural mutations in IFITM3 modulate post□translational regulation and toggle antiviral specificity. EMBO Rep. 17, 1657–1671 (2016).

