## Supplementary material for "Genomic and transcriptomic characterization of Delta SARS-CoV-2 infection in free-ranging white-tailed deer (*Odocoileus virginianus*)": Figure S1, Figure S2, Figure S3, Figure S4, Figure S5

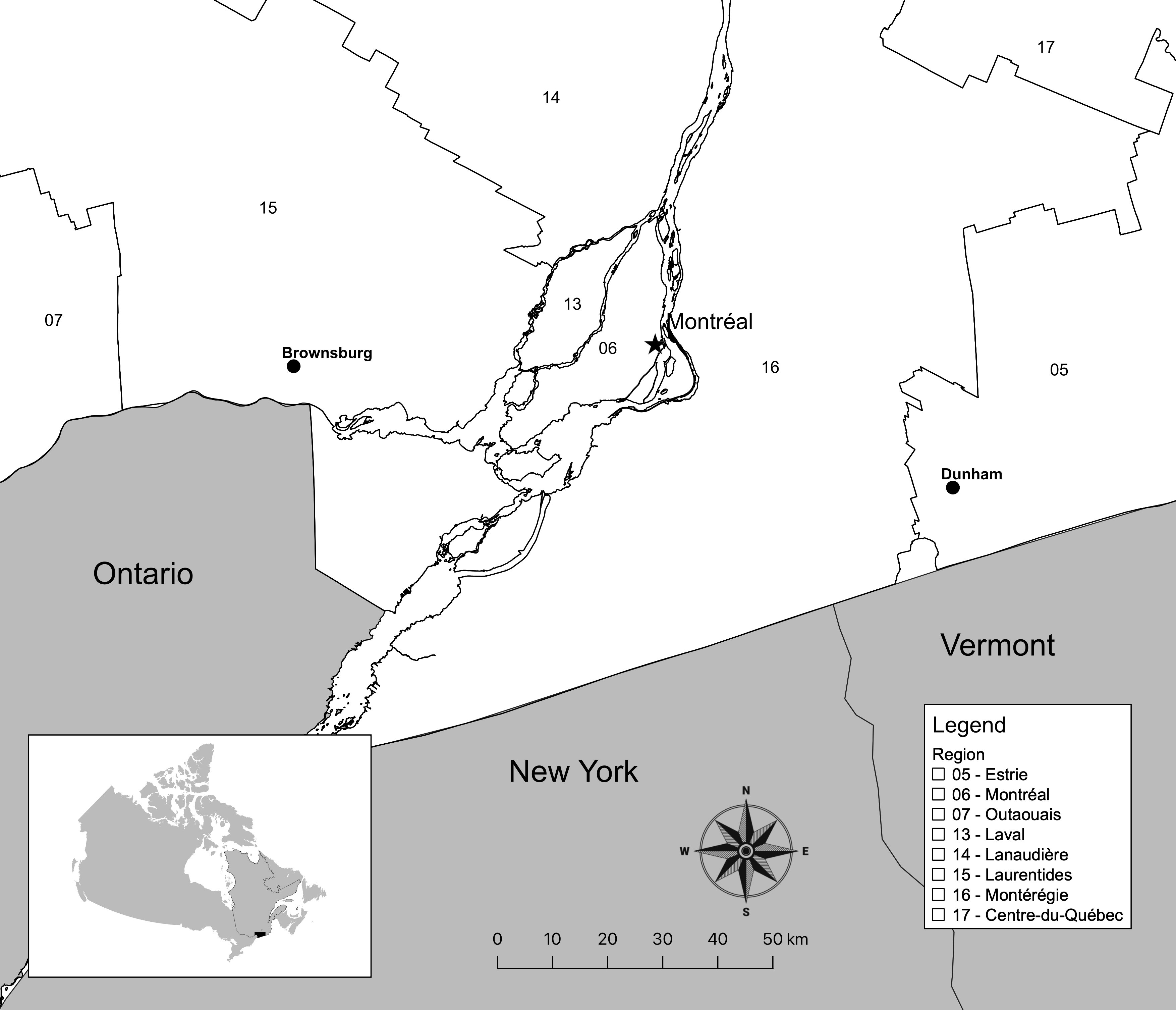

**Figure S1.** Map of southern Québec administrative regions and corresponding identification numbers within the study region. Inset shows location of Québec (outlined) and study region (shaded black) within Canada.

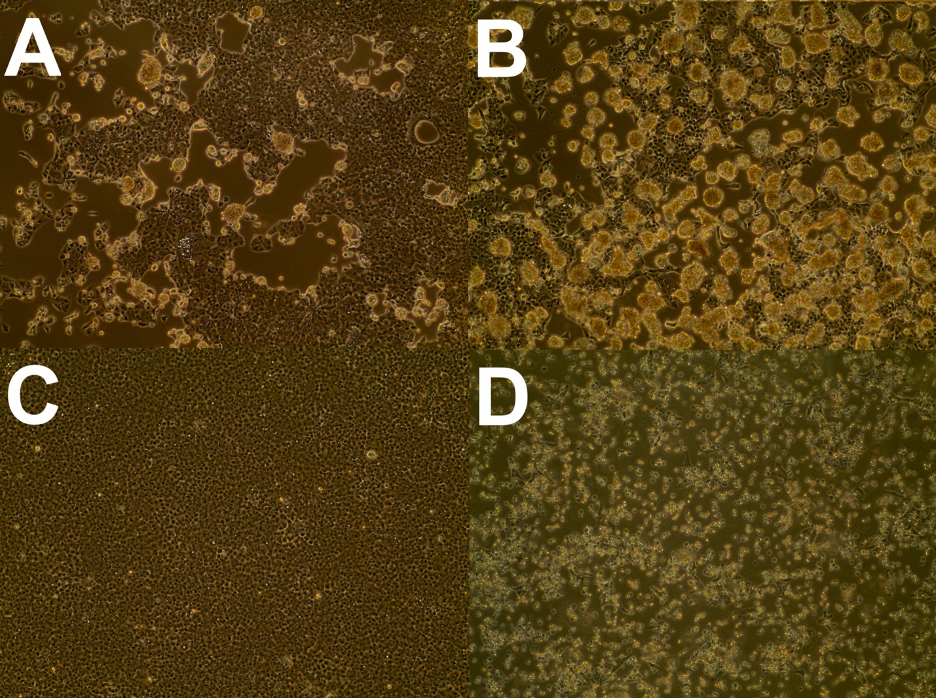

**Figure S2.** Observed cytopathic effect (CPE) in VeroE6 cells inoculated with nasal swabs from SARS-CoV-2 positive white-tailed deer at 5 days post infection (A-4055, B-4249) with a mock inoculated negative control (C) and positive control inoculated with a nasopharyngeal clinical specimen from a COVID-19 patient (D). Magnification was 100x.

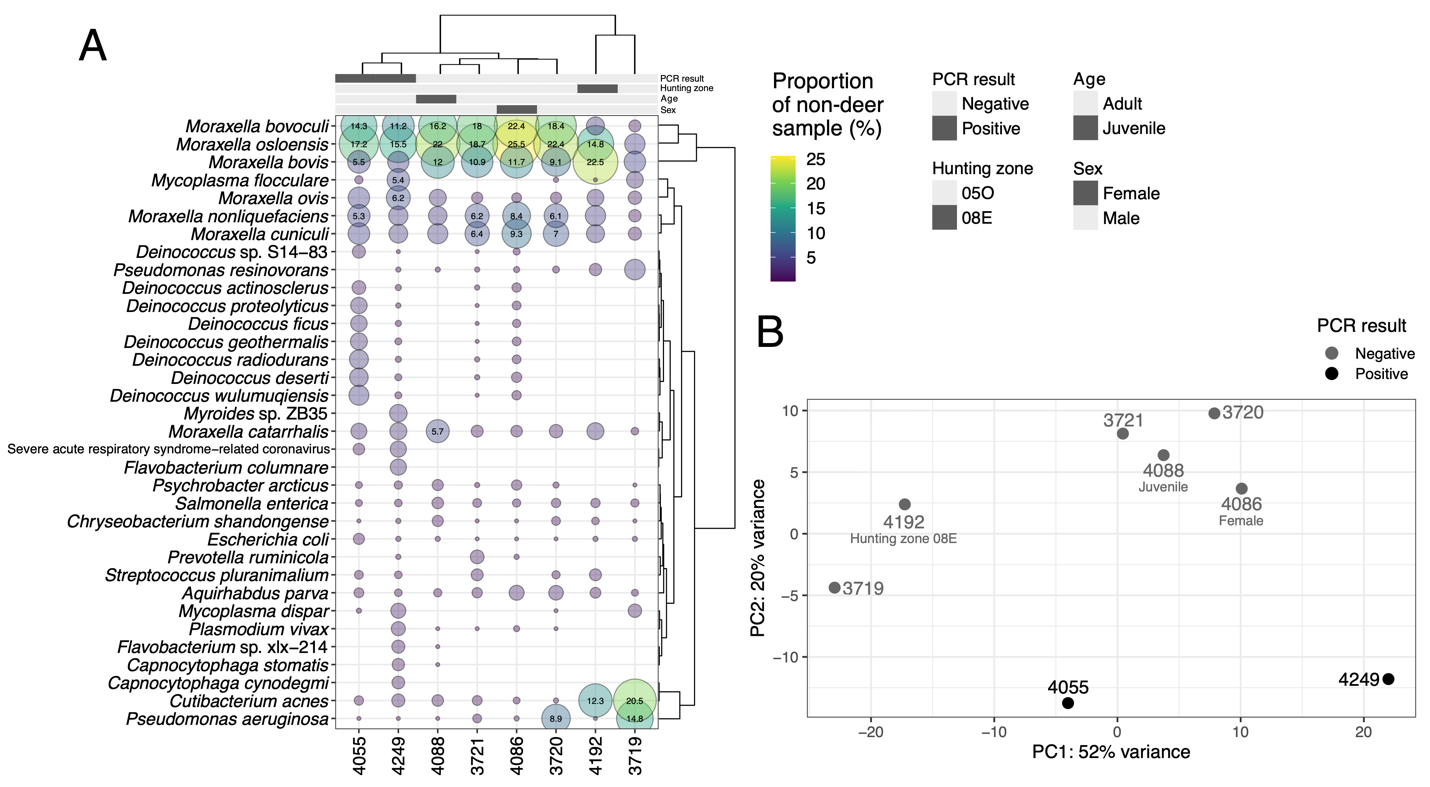

**Figure S3.** RNA-seq taxonomic profiling and PCA of deer transcriptome expression profile. A) Top community members after removal of deer reads. Proportion numbers are only displayed if larger than 5.0%. PCR result refers to the SARS-CoV-2 RT-PCR analysis (see Methods). B) An ordination of the overall WTD gene expression profile. PCA created with the VST transformed gene expression profiles. Metadata other than the PCR result is displayed on the graph under the sample ID.

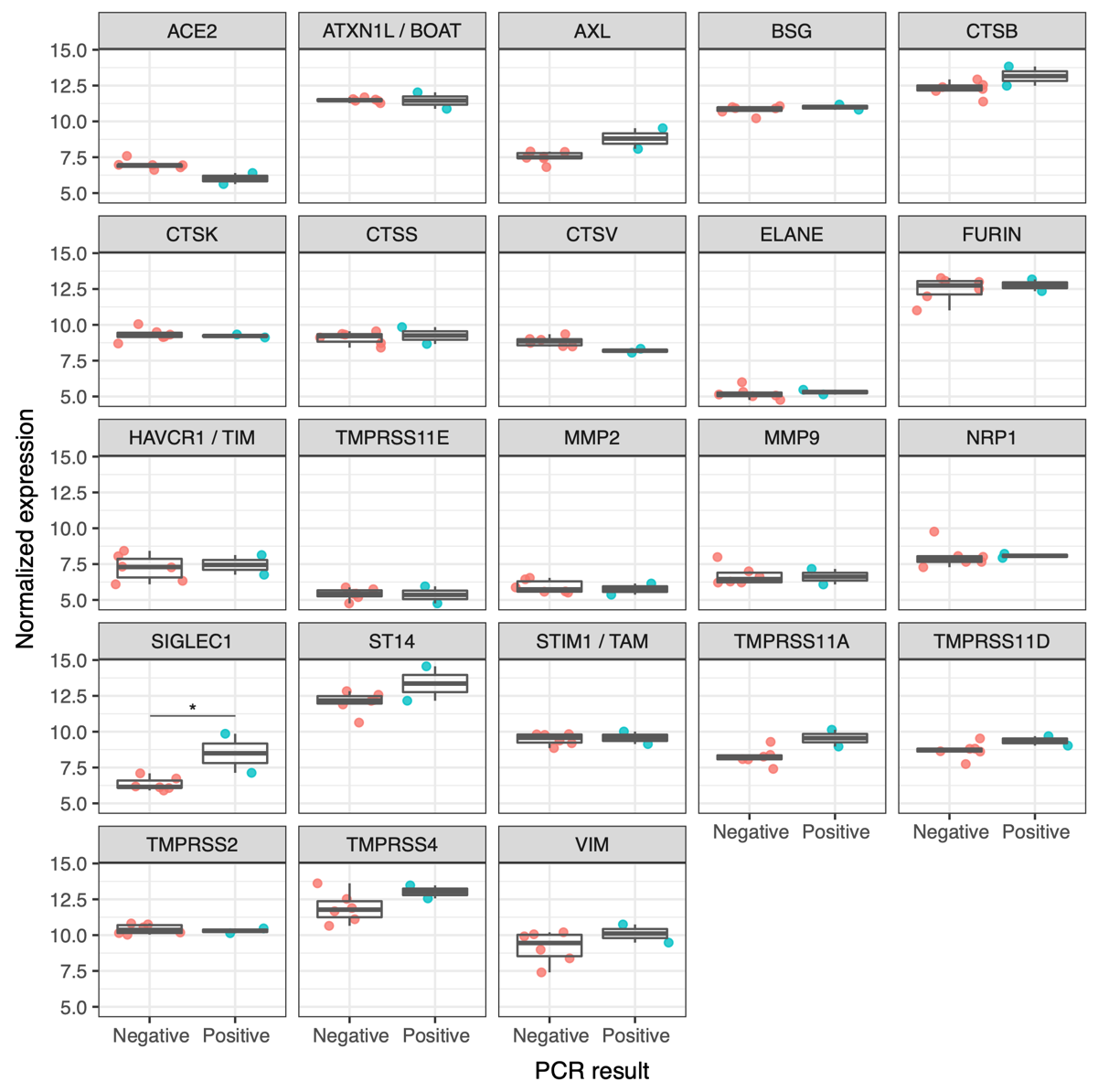

**Figure S4.** Select SARS-CoV-2 viral attachment and entry factors important in human COVID-19 infections with their VST transformed expression value displayed for each deer sample. Genes are labelled with their inferred human gene ortholog. The PCR result identifies which deer samples were SARS-CoV-2 RT-PCR-positive or -negative. Significant differentially expressed genes (adjusted p-value<0.05) are indicated with an asterisk (*).

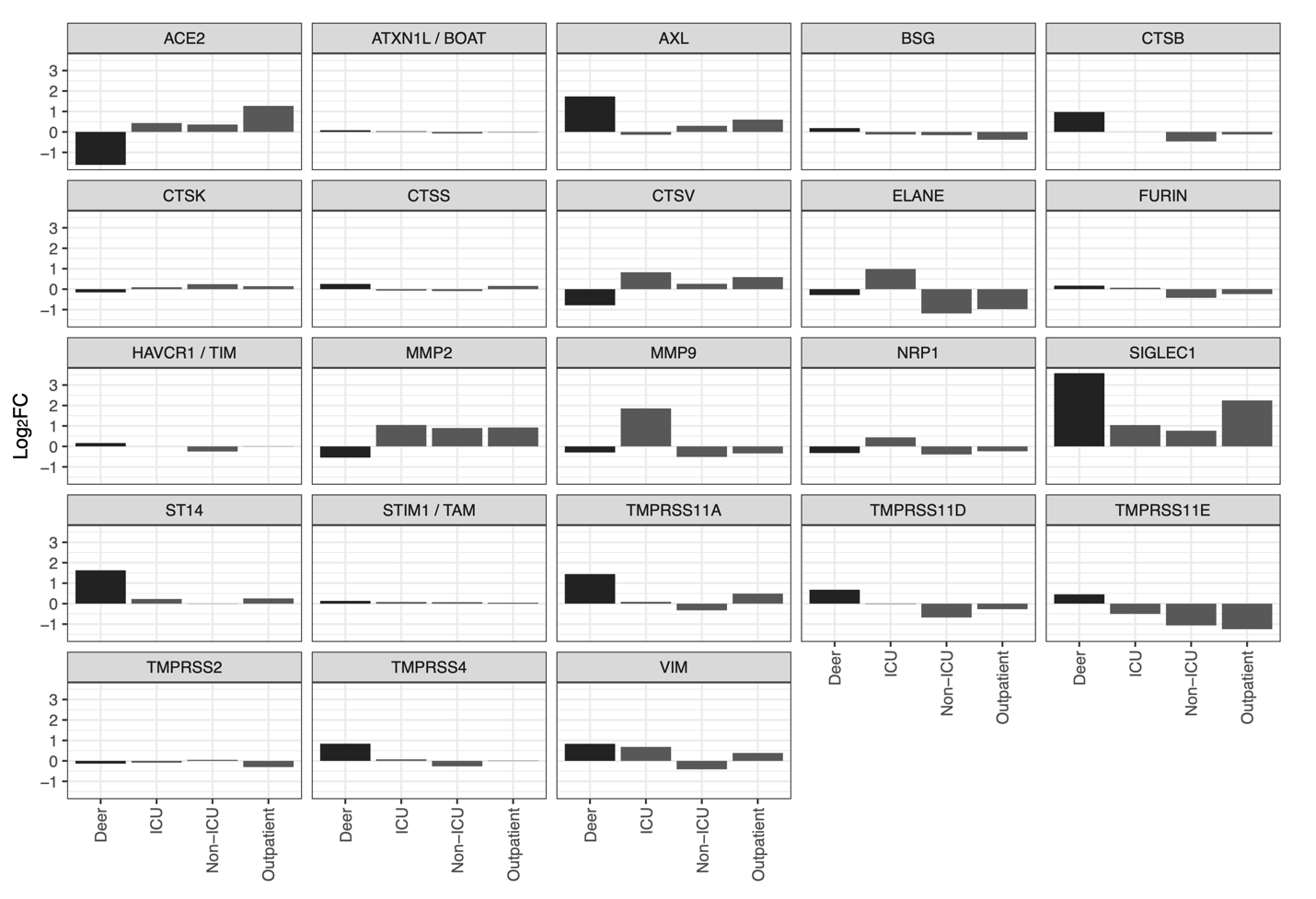

**Figure S5.** Select SARS-CoV-2 viral attachment and entry factors important in human COVID-19 infections with their log_2_ fold change values in deer versus human COVID-19 infections. Human nasopharyngeal swab samples in ICU, non-ICU, and outpatient individuals were compared to nasopharyngeal swab samples from individuals without SARS-CoV-2. Those log_2_ fold change values are displayed here against the deer nasal swab differential gene expression analysis log_2_ fold change values. *HAVCR1* here corresponds to the WTD gene *LOC110145616*. The genes in this figure are labelled with their human gene IDs.
