## supplemental methods for "Genomic and transcriptomic characterization of Delta SARS-CoV-2 infection in free-ranging white-tailed deer (*Odocoileus virginianus*)"

*SARS-CoV-2 amplification and sequencing*

SARS-CoV-2 whole genome sequencing was performed at Sunnybrook Research Institute (SRI) and analyzed at CFIA. Extracted RNA was reverse transcribed into cDNA; 8 μL RNA was mixed with 8 μL nuclease-free water and 4 μL LunaScript RT SuperMix (New England BioLabs) then incubated at 25°C for 2 minutes followed by an incubation at 55°C for 20 minutes and 95°C for 1 minute before holding at 4°C.

cDNA was amplified using ARTIC v3 primer pools (https://github.com/artic-network/artic-ncov2019). Two separate reactions were prepared by combining 2.5 μL cDNA with 6 μL nuclease-free water, 12.5 μL Q5 Hot Start High-Fidelity 2X Master Mix (New England BioLabs) and 4 μL of 10 μM primer pool 1 or primer pool 2 (Integrated DNA Technologies; https://www.idtdna.com). The PCR cycling conditions included an initial denaturation at 98°C for 30 seconds followed by 35 cycles of 98°C for 15 seconds and 63°C for 5 minutes before holding at 4°C. Both reactions were combined and purified with 1X ratio Sample Purification Beads (Illumina; https://www.illumina.com). The resulting amplicons were quantified using Qubit 1X dsDNA HS Assay Kit (Thermo Fisher Scientific).

Libraries were constructed using Illumina DNA Prep (Illumina) and IDT for Illumina DNA/RNA UD Indexes (Illumina) and sequenced on Illumina MiniSeq using 2x149 paired-end reads. Six negative controls (i.e., two extraction, two RT-PCR, and two library prep controls) are included in each sequencing run and all controls must contain <1% of SARS-CoV-2 genome for the run to pass quality control.

Paired-end Illumina reads for samples 4055, 4204, 4205 and 4249 were analyzed using the nf-core/viralrecon Nextflow workflow (v2.2) (Ewels et al., 2020; Patel et al., 2021; Di Tommaso et al., 2017), which performed the following analysis steps: FastQC (v0.11.9) read quality assessment (https://www.bioinformatics.babraham.ac.uk/projects/fastqc/); fastp (v0.20.1) read quality filtering and trimming (Chen et al., 2018); read mapping to Wuhan-Hu-1 (MN908947.3) SARS-CoV-2 reference sequence with Bowtie2 (v2.4.2) (Langmead and Salzberg, 2012); read mapping statistics calculation with Mosdepth (v0.3.1) (Pedersen and Quinlan, 2017) and Samtools (v1.12) (Danecek et al., 2021; Li et al., 2009); ARTIC V3 primer trimming, variant calling and consensus sequence generation with iVar (v1.3.1) (Grubaugh et al., 2019); variant effect analysis and summarization with SnpEff (v5.0) (Cingolani et al., 2012a) and SnpSift (v4.3t) (Cingolani et al., 2012b), respectively; SARS-CoV-2 lineage assignment with Pangolin (v3.1.17).

**References:**

Chen, S., Zhou, Y., Chen, Y., Gu, J., 2018. fastp: an ultra-fast all-in-one FASTQ preprocessor. Bioinformatics 34, i884–i890. https://doi.org/10.1093/bioinformatics/bty560

Cingolani, P., Patel, V.M., Coon, M., Nguyen, T., Land, S.J., Ruden, D.M., Lu, X., 2012a. Using Drosophila melanogaster as a Model for Genotoxic Chemical Mutational Studies with a New Program, SnpSift. Front Genet 3, 35. https://doi.org/10.3389/fgene.2012.00035

Cingolani, P., Platts, A., Wang, L.L., Coon, M., Nguyen, T., Wang, L., Land, S.J., Lu, X., Ruden, D.M., 2012b. A program for annotating and predicting the effects of single nucleotide polymorphisms, SnpEff. Fly 6, 80–92. https://doi.org/10.4161/fly.19695

Danecek, P., Bonfield, J.K., Liddle, J., Marshall, J., Ohan, V., Pollard, M.O., Whitwham, A., Keane, T., McCarthy, S.A., Davies, R.M., Li, H., 2021. Twelve years of SAMtools and BCFtools. GigaScience 10, giab008. https://doi.org/10.1093/gigascience/giab008

Di Tommaso, P., Chatzou, M., Floden, E.W., Barja, P.P., Palumbo, E., Notredame, C., 2017. Nextflow enables reproducible computational workflows. Nat Biotechnol 35, 316–319. https://doi.org/10.1038/nbt.3820

Ewels, P.A., Peltzer, A., Fillinger, S., Patel, H., Alneberg, J., Wilm, A., Garcia, M.U., Di Tommaso, P., Nahnsen, S., 2020. The nf-core framework for community-curated bioinformatics pipelines. Nat Biotechnol 38, 276–278. https://doi.org/10.1038/s41587-020-0439-x

Grubaugh, N.D., Gangavarapu, K., Quick, J., Matteson, N.L., De Jesus, J.G., Main, B.J., Tan, A.L., Paul, L.M., Brackney, D.E., Grewal, S., Gurfield, N., Van Rompay, K.K.A., Isern, S., Michael, S.F., Coffey, L.L., Loman, N.J., Andersen, K.G., 2019. An amplicon-based sequencing framework for accurately measuring intrahost virus diversity using PrimalSeq and iVar. Genome Biology 20, 8. https://doi.org/10.1186/s13059-018-1618-7

Langmead, B., Salzberg, S.L., 2012. Fast gapped-read alignment with Bowtie 2. Nat Methods 9, 357–359. https://doi.org/10.1038/nmeth.1923

Patel, H., Varona, S., Monzón, S., Espinosa-Carrasco, J., Heuer, M.L., Nf-Core Bot, Underwood, A., Gabernet, G., Ewels, P., MiguelJulia, Kelly, S., Stevin Wilson, Erika, Sameith, K., Garcia, M.U., Jcurado, Menden, K., 2022. nf-core/viralrecon: nf-core/viralrecon v2.5 - Manganese Monkey. https://doi.org/10.5281/ZENODO.3901628

Pedersen, B.S., Quinlan, A.R., 2018. Mosdepth: quick coverage calculation for genomes and exomes. Bioinformatics 34, 867–868. https://doi.org/10.1093/bioinformatics/btx699
